# Small extracellular vesicles from young mice prevent frailty, improve healthspan and decrease epigenetic age in old mice

**DOI:** 10.1101/2021.07.29.454302

**Authors:** Jorge Sanz-Ros, Cristina Mas-Bargues, Daniel Monleón, Juozas Gordevicius, Robert T. Brooke, Mar Dromant, Aksinya Derevyanko, Ana Guío-Carrión, Aurora Román-Domínguez, Nekane Romero-García, Marta Inglés, María A. Blasco, Steve Horvath, Jose Viña, Consuelo Borrás

## Abstract

Aging is associated with an increased risk of frailty, disability, comorbidities, institutionalization, falls, fractures, hospitalization, and mortality. Searching for strategies to delay the degenerative changes associated with aging and frailty is interesting. We treated old animals intravenously with small extracellular vesicles (sEVs) derived from adipose mesenchymal stem cells (ADSCs) of young animals, and we found an improvement of several functional parameters usually altered with aging, such as motor coordination, grip strength, fatigue resistance, fur regeneration, and renal function. Frailty index analysis showed that 40% of old control mice were frail, whereas none of the old ADSCs-sEVs treated mice were. Molecular and structural benefits in muscle and kidney accompanied this functional improvement. ADSCs-sEVs induced pro-regenerative effects and a decrease in oxidative stress, inflammation, and senescence markers. Moreover, predicted epigenetic age was lower in tissues of old mice treated with ADSCs-sEVs and their metabolome changed to a youth-like pattern. Finally, we gained some insight into the miRNAs contained in sEVs that might be, at least in part, responsible for the effects observed. We propose that young sEVs treatment can be beneficial against frailty and therefore can promote healthy aging.

## Introduction

To add health to the years gained, as well as to promote the ability to live autonomously is a public health priority in modern societies. The search for strategies to delay the degenerative changes associated with aging and frailty is particularly interesting.

Aging is accompanied by an impairment in the physical condition and an increased risk of frailty^1^. It is characterized by several changes at the cellular and organismal levels that result in a decreased functionality of several tissues. Alterations in intercellular communication have been described as potential drivers of age-related dysfunctions^2^. Cellular function not only depends on cell-autonomous factors, but is also affected by the extracellular environment, and its modification can have a great effect on the performance of several tissues^3, 4^. Parabiosis experiments conducted in mice demonstrated that factors present in the blood of a young organism are beneficial for an aged one, improving several parameters affected by aging^5,6,7^.

Extracellular vesicles (EVs), small vesicles that are released by virtually all cell types, with an innate ability to mediate the transmission of signalling molecules (proteins, small RNAs, DNA) between cells are among the factors that are involved in the communication between cells^8^.

Allogenic transplantation of mesenchymal stem cells in frail people has shown very promising results to treat frailty^9^. Stem cells have intrinsic regenerative effects that are not only mediated by the repopulation of damaged tissue. The releasing of regulatory molecules is also proposed as one of the most important mechanisms in stem cell therapies^10, 11^. More specifically, sEVs derived from multiple stem cells have demonstrated their capacity to promote tissue regeneration after several types of damage^12, 13^. Compared to stem cells, sEVs are more stable, have no risk of aneuploidy, have a lower chance of immune rejection, and can provide an alternative therapy for various diseases^14,15,16,17^.

It is important to consider that the way of culturing stem cells affects dramatically paracrine signals. We previously showed that oxygen tension in culture is one of the most important factors, affecting the biological function of sEVs released by these cells^18,19,20^, and that sEVs derived from mesenchymal stem cells cultured under hypoxic conditions can mediate the transfer of miRNAs to senescent cells and improve cellular function^19^. A recent study demonstrated that hypothalamic stem cells modulate aging speed through the release of miRNAs contained in EVs^21^. sEVs are emerging as a potential therapy in the aging field. It has been recently shown that sEVs can exert pro-regenerative effects in tissues of old mice, as well as a decrease in senescence-related damage^22,23,24,25^. However, the effect of mesenchymal stem cells-derived sEVs on the healthspan of an aged organism has not been fully addressed. Here we show that sEVs from young adipose-derived stem cells (ADSCs-sEVs) improve several functions that are impaired in old mice. Old mice that received young ADSCs-sEVs were protected against frailty and improved on physical condition tests, fur regeneration, and renal function. ADSCs-sEVs induced pro-regenerative effects on muscle and kidney of aged mice, as well as a decrease in oxidative stress, inflammation, and senescence markers. Moreover, predicted epigenetic age was lower in tissues of old mice treated with ADSCs-sEVs and the metabolome of old mice treated with ADSCs-sEVs changed from an old-like pattern to a youth-like one. Finally, we gained some insight into the miRNAs contained in sEVs that might be, at least in part, responsible for the effects observed.

## Results

### ADSCs-sEVs improve healthspan and prevent frailty in old mice

As aged organisms typically show impairment in their physical condition and an increased risk of frailty, we designed an experiment to test if ADSCs-sEVs could have a beneficial effect on the physical performance of old mice. Before treatment, we measured total body weight, grip strength, motor coordination, and resistance to fatigue in aged mice, which were recorded as baseline values. Twenty-four hours later, each mouse received an injection of ADSCs-sEVs or PBS as a control (day 1). One week later (day 7), mice received a second injection (as outlined in Figure 1a).

**Figure 1:**
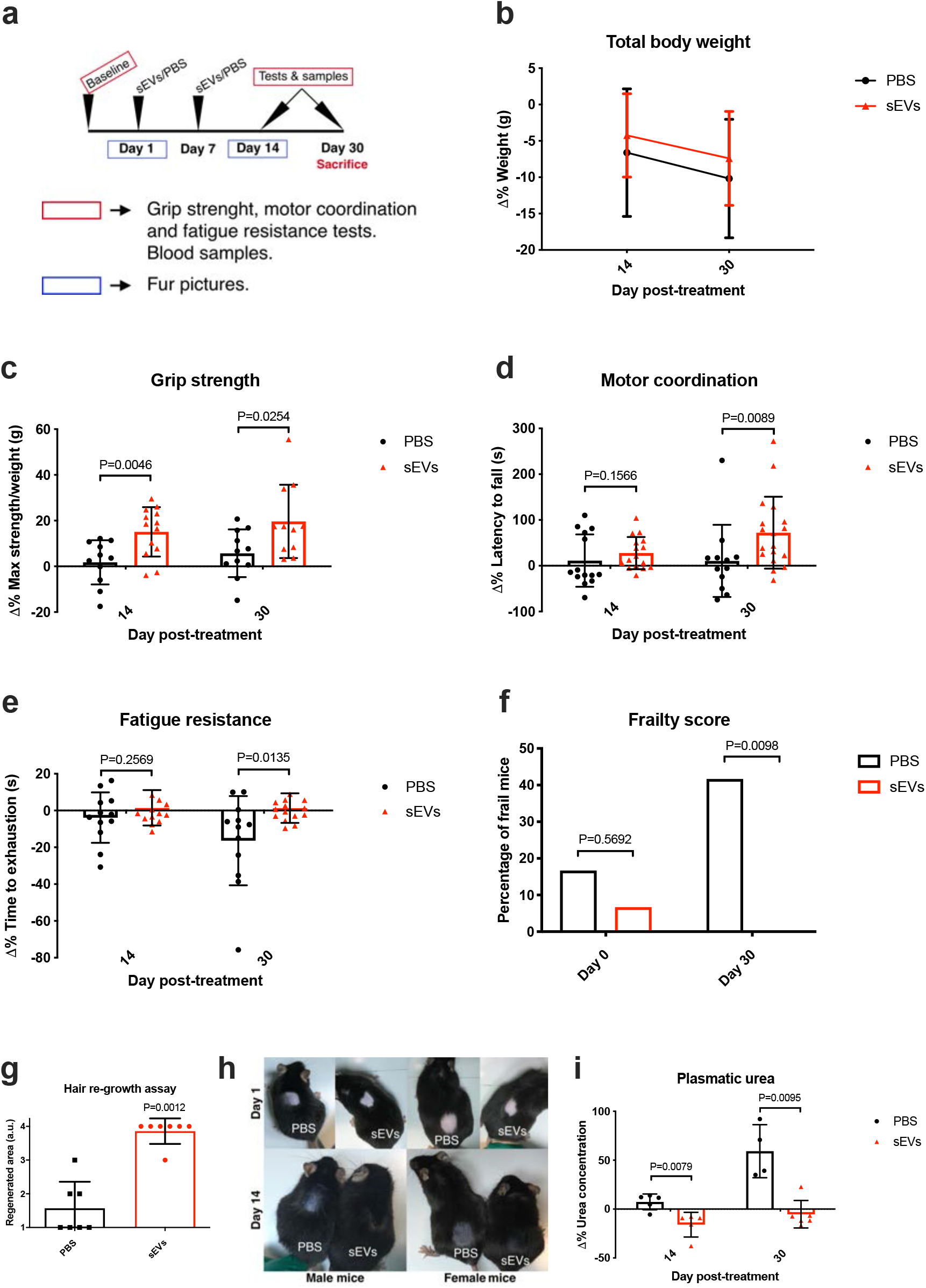
ADSCs-sEVs improve healthspan and prevent frailty in old mice. a: Schematic representation of the experiment design. b: Quantification of the change in total body weight and physical condition tests, data is shown as the decrease in percentage from baseline before treatment with sEVs/PBS. PBS day 14 n=12, day 30 n=12, sEVs day 14 n=14, day 30 n=15. c-e: Quantification of the change in physical condition tests, data is shown as the increase in percentage from baseline before treatment with sEVs/PBS. Grip strength: PBS day 14 n=11, day 30 n=11, sEVs day 14 n=13, day 30 n=11. Motor coordination: PBS day 14 n=14, day 30 n=12, sEVs day 14 n=16, day 30 n=18. Fatigue resistance: PBS day 14 n=12, day 30 n=12, sEVs day 14 n=14, day 30 n=15. f: Quantification of the number of frail mice before and 30 days after treatment with sEVs/PBS, determined by a score based on the clinical phenotype of frailty. PBS day 0 n=12, day 30 n=12, sEVs day 0 n=15, day 30 n=15. g: Quantification n of the hair re-growth capacity of dorsal skin in old mice. PBS n=7, sEVs n=7. h: Representative image of hair regeneration in both male and female mice before and 14 days after treatment with sEVs/PBS. i: Quantification of the changes in plasmatic urea, data is shown as the increase in percentage from basal levels of urea in plasma before treatment with sEVs/PBS. PBS day 14 n=5, day 30 n=4, sEVs day 14 n=5, day 30 n=6. All data are shown as mean±SD.

As physical tests depend on the weight of mice, we measured the total body weight of mice during the experiment, and we did not find differences between groups. In addition, we failed to observe any signs of toxicity (excessive weight loss) during the entire experiment (Figure 1b). Physical tests were repeated on days 14 and 30 after ADSCs-sEVs or PBS treatment. On day 14 we could observe an improvement in the strength test on ADSCs-sEVs treated mice. The maximum benefit was observed on day 30: mice treated with ADSCs-sEVs showed an improvement in grip strength, motor coordination, and fatigue resistance when compared to controls (Figure 1c-e).

Interestingly, 60 days after treatment, the positive effect was lost, and we could not find any differences between groups (Supplementary Table 1). Thus, the protective effects of ADSCs-sEVs are, as expected, transitory.

For a quantitative assessment of frailty, we used a score based on the clinical phenotype of frail humans developed by our group ^26^. This enabled us to classify each mouse as frail or non-frail at each time point. On day 30, there were no frail mice in the ADSCs-sEVs treated group, while the control group shed 40% of frail mice, which is in accordance with previous results in mice of this age^26^ (Figure 1f). Fur density is reduced and capacity to re-growth hair becomes impaired with aging^27^. To check the effect of ADSCs-sEVs on the growth of hair, we plucked a square of 1cm x 1cm of the dorsal fur of each mouse on day 1 just before injection. On day 14, we observed that most mice treated with ADSCs-sEVs had regenerated the entire area. In contrast, mice injected with PBS had a much lower capacity for hair re-growth (Figure 1g-h).

To determine possible changes in the renal function we used plasma values of urea, as it is a well-known marker of kidney function. We obtained plasma samples of each mouse on days 0, 14, 30, and 60 and measured urea levels at each time point. Day 0 levels were used as a baseline. Fourteen days after being injected with ADSCs-sEVs, mice showed a decrease in plasma concentration of urea, whereas the control group showed no modification. On day 30, urea levels raised 50% from baseline in the control group. By contrast, the ADSCs-sEVs group showed a plasmatic urea level like day 0 (Figure 1i).

In addition, as a control to further test the possible effect of age in the quality of sEVs and ADSCs, we performed a pilot study with 3 old mice treated with ADSCs-sEVs isolated from old mice, we did not find any effect of these old-sEVs in physical performance tests, indicating that the age of ADSCs donor is essential in the beneficial effects observed (Supplementary Table 2).

Taken together, we find that only ADSCs-sEVs isolated from young mice induce an improvement in the global healthspan of aged mice.

### ADSCs-sEVs reverse age-related structural changes in kidney and muscle of old mice

As an organism begins to age, its tissues suffer from several structural changes, which are detrimental to the normal function of the tissue, such as a loss of regenerative capacity and fibrosis^2, 28^. We selected kidney and muscle to test the effect of ADSCs-sEVs on tissue structure, as these organs are specially involved in the aging process: kidney function is one of the organs that age faster in normal individuals^29^, and muscle function is essential for the activities of daily life and is directly related to frailty^30^. Therefore, we obtained kidney and muscle from old mice 30 days after treatment with ADSCs-sEVs/PBS. In the kidney, we performed histological analysis of the renal cortex, looking for the presence of tubular atrophy and interstitial fibrosis, two distinctive changes associated with aging that leads to the loss of renal function^31^. The macroscopic appearance of kidneys was very different between old untreated and treated mice, kidneys isolated from old mice treated with sEVs were more like those isolated from young mice (Figure 2a). Control mice showed widespread tubular atrophy, defined by a reduced tubular density and the presence of dilated tubules. In contrast, tubular density was much higher in mice treated with ADSCs-sEVs, as well as very low tubular dilation. We did not observe differences in glomerular density between groups (Figure 2b-e). For the study of interstitial fibrosis, we used Sirius red staining to measure collagen deposition. We noted a mild decrease in the ADSCs-sEVs treated group, although no statistically significant differences were observed (Figure 2f).

**Figure 2:**
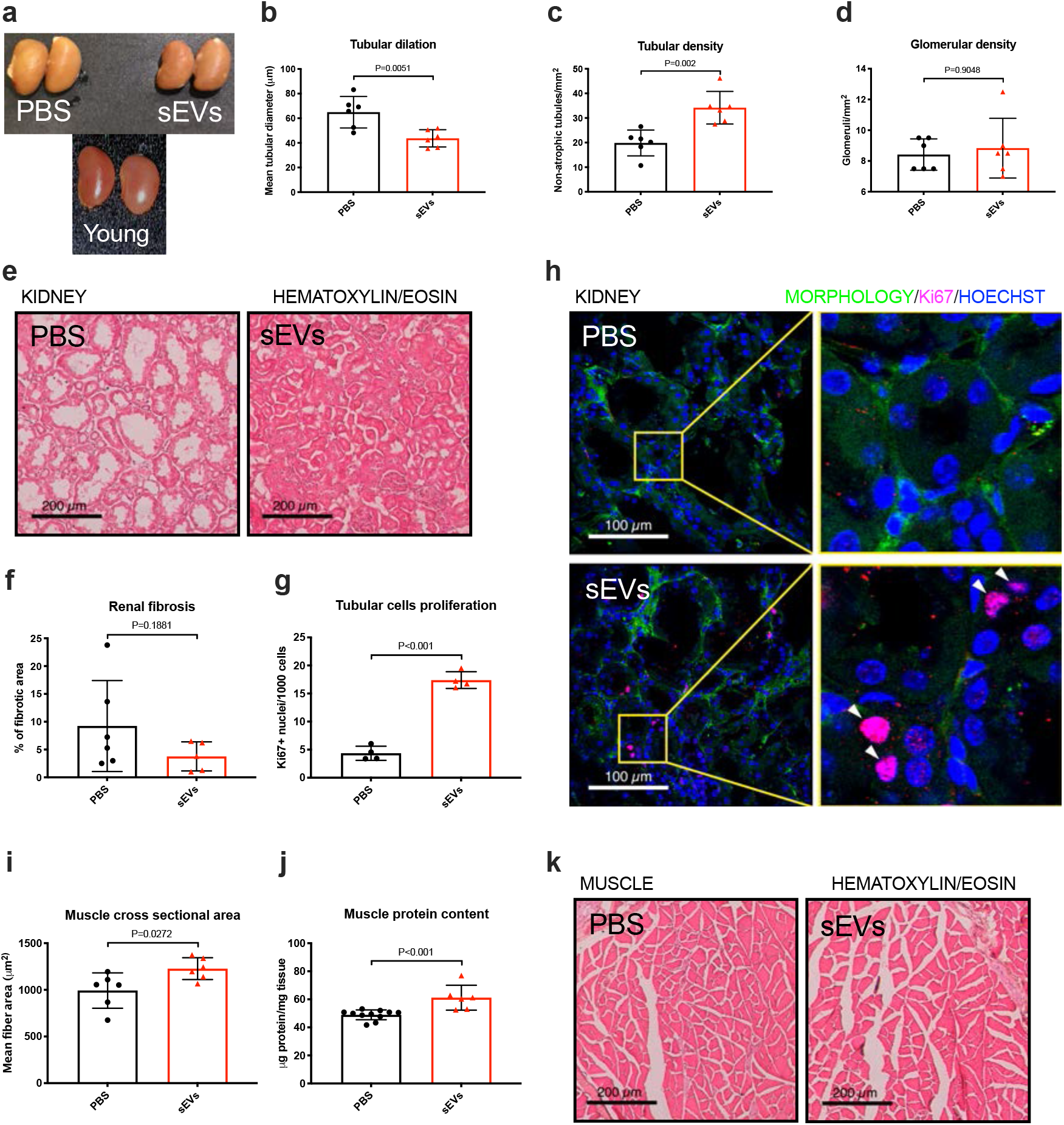
ADSCs-sEVs induce pro-regenerative effects on kidney and muscle of old mice. a: Macroscopic image of kidneys isolated from old mice treated with PBS/sEVs. b-d: Quantification of histological changes in the kidney (renal cortex) from old mice induced by sEVs. PBS n=6, sEVs n=6. e: Haematoxylin-Eosin representative images of the effects on the structure of the renal cortex of old mice. f: Quantification of extracellular fibrosis by Sirius red staining. PBS n=6, sEVs n=6. g: Quantification of Ki67+ cells in the renal tubules of old mice. PBS n=4, sEVs n=4. h: Representative immunofluorescence images showing Ki67+ cells in the renal tubules of old mice. i-j: Quantification of the mean cross-sectional area (CSA) of muscle fibers and total protein content in the gastrocnemius of old mice. CSA: PBS n=6, sEVs n=6. Protein content: PBS n=11, sEVs n=6. k: Haematoxylin-Eosin representative images of gastrocnemius of old mice. All data are shown as mean±SD.

These observations suggested that ADSCs-sEVs may induce the proliferation of tubular cells as a mean of regeneration, as these cells can repopulate renal tubules after damage^32^. Using Ki67 as a proliferation marker, we looked for the presence of Ki67+ cells in the cortex tubules of our mice. We found very low levels of proliferating cells in the control group, however, in the treated group we identified a clear population of Ki67+ cells in the tubules, indicating active proliferation of the tubular cells in these mice (Figure 2g-h).

Muscle loss of function and atrophy lead to a global dysfunction of the organism as it ages, as the musculoskeletal system is essential for the activities of daily life. The prevalence of sarcopenia (defined by a loss of muscle mass and function) increases with age^33^. To investigate the effect of ADSCs-sEVs on muscle tissue of old mice, we measured the cross-sectional area (CSA) of muscle fibers, which is a parameter related to muscle atrophy and loss of strength. Interestingly, we found an increase in the mean CSA with ADSCs-sEVs treatment, which was supported by a higher protein concentration in the muscles of ADSCs-sEVs treated mice (Figure 2i-k).

In sum, these results show that ADSCs-sEVs induce pro-regenerative effects in kidney and muscle, partially reverting structural changes associated with aging in these tissues.

### ADSCs-sEVs mitigate molecular traits associated with aging in kidney and muscle and lower senescence in old mice and in an *in vitro* model

Multiple molecular traits contribute to the aging process^2^. Here we have studied the effect of ADSCs-sEVs treatment on several markers associated with aging: oxidative stress, senescence, inflammation, and telomere attrition. We obtained kidneys and muscle from old mice 30 days after treatment with ADSCs-sEVs /PBS and studied how the treatment affected the presence of these markers. In the case of oxidative stress, we examined the levels of oxidation in two macromolecules, lipids, and proteins. We used malondialdehyde (MDA) as a marker of lipid peroxidation, and we observed lower levels of MDA in the muscle of mice treated with ADSCs-sEVs, with no changes in the kidney (Figure 3a). In protein oxidation levels, determined by carbonyl groups present in oxidized proteins, we found similar results as with those of MDA, with lower oxidation in the muscle of mice treated with ADSCs-sEVs, and no changes in the kidney (Figure 3b).

**Figure 3:**
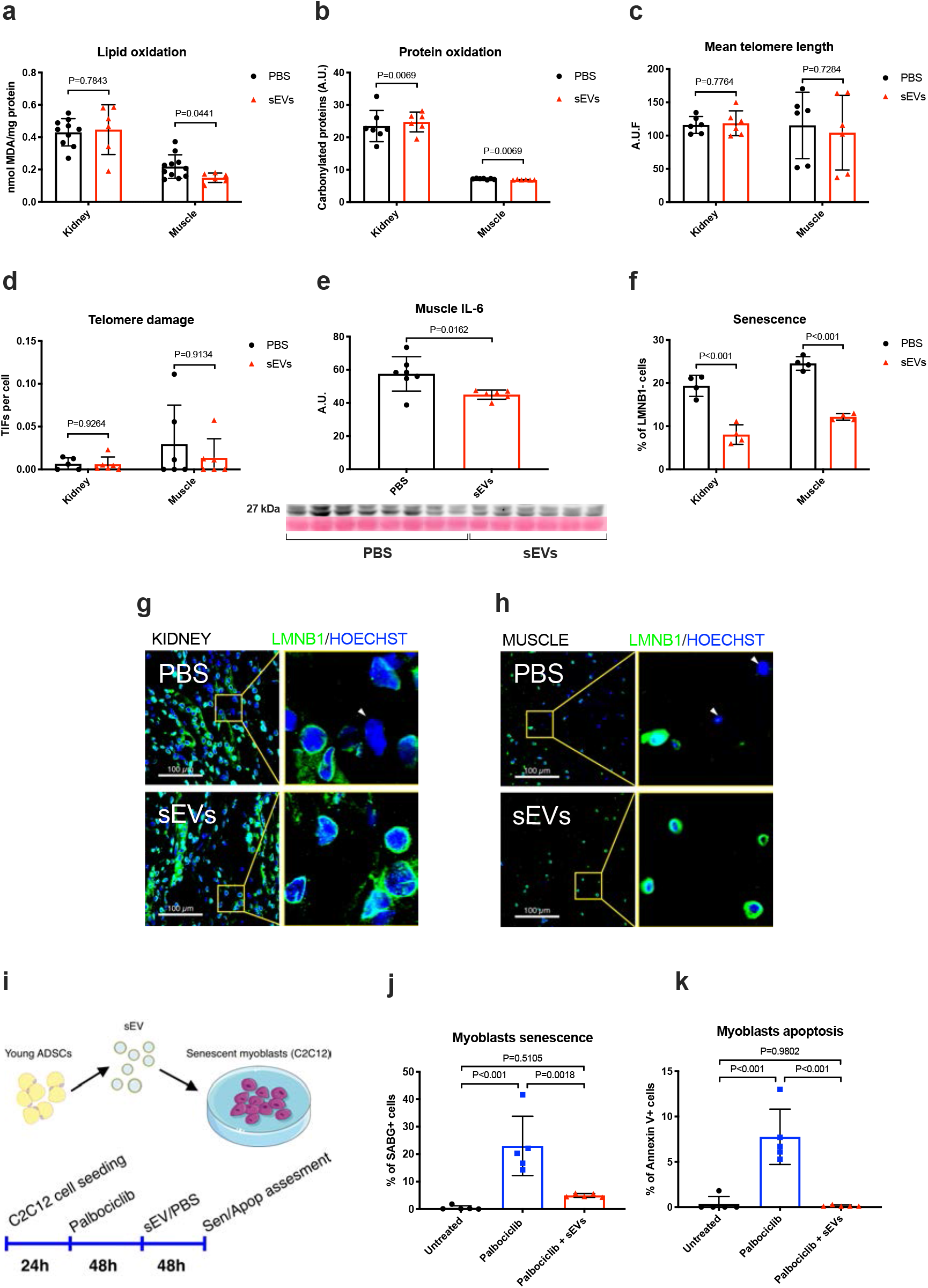
sEVs from young ADSCs ameliorate molecular traits of aging in kidney and muscle and lower senescence in old mice and in an *in vitro* model. a: Quantification of malondialdehyde as a marker of lipid peroxidation in kidney and muscle from old mice. PBS kidney n=10, muscle n=11, sEVs kidney n=6, muscle n=6. b: Quantification of protein carbonylation as a marker of protein oxidation in kidney and muscle of old mice. PBS kidney n=7, muscle n=7, sEVs kidney n=6, muscle n=6. c: Mean telomere length measured as arbitrary units of fluorescence (a.u.f) in kidney and muscle of old mice. PBS kidney n=6, muscle n=6, sEVs kidney n=6, muscle n=6. d: Quantification of telomere dysfunction-induced foci (TIF) measured as number of telomeric probe and DNA damage marker 53BP1 colocalizing foci per cell by Immuno FISH in kidney and muscle of old mice. PBS kidney n=5, muscle n=5, sEVs kidney n=6, muscle n=6. e: Quantification of IL-6 in the muscle of old mice. PBS n=7, sEVs n=6. f-h: Quantification and representative immunofluorescence images of LMNB1 loss in gastrocnemius of old mice. PBS kidney n=4, muscle n=4, sEVs kidney n=4, muscle n=4. i: Graphical representation of *in vitro* experiment with senescent murine myoblasts. j-k: Assessment of senescence-associated β-galactosidase activity and Annexin V by flow cytometry in senescent murine myoblasts. Untreated n=5, Palbociclib n=5, Palbociclib + sEVs n=5. All data are shown as mean±SD.

Telomere attrition and damage to telomeres are two of the most studied factors that lead to genomic instability and loss of proliferative capacity. Telomere shortening is observed in many species during normal aging, and excessively short telomeres are associated with multiple conditions characterized by a loss of regenerative capacity, such as pulmonary fibrosis, dyskeratosis congenita, and aplastic anaemia^34^. We explored the effect of ADSCs-sEVs on telomere length and damage, as possible mediators of the observed effects in mice. For this purpose, we measured mean telomere length and DNA damage at the telomeres quantified as telomere dysfunction induced foci (TIF). We did not find significant differences between both groups (Figure 3c-d), although we found that the muscle of treated mice showed slightly lower telomere damage than the non-treated ones.

Senescent cells accumulate in aged tissues, having a detrimental effect on their function, as they are unable to enter the cell cycle. In addition, these cells act in a pleiotropic manner releasing several factors that make up the senescence-associated secretory phenotype (SASP), contributing to the development of age-associated diseases^35, 36^. We measured the loss of lamin B1 (LMNB1), a robust senescence marker^37^, in muscle and kidney of old mice 30 days after treatment with ADSCs-sEVs/PBS and noticed an important decrease in the number of senescent cells in both tissues in mice treated with ADSCs-sEVs (Figure 3f-h).

Chronic inflammation is a condition closely related to aging and senescence, an increase in pro-inflammatory cytokines in several tissues has been linked to aging, frailty, and age-related diseases^38^. IL-6 is one of the most important factors of the SASP, so we measured levels of this interleukin in the muscle of old mice, observing a decrease in its levels in muscle of ADSCs-sEVs treated mice (Figure 3e).

Finally, to test the effect of ADSCs-sEVs on a more controlled environment, we developed an *in vitro* model of senescence in muscle progenitor cells. For this purpose, we used C2C12 mouse myoblasts and induced cellular senescence with Palbociclib (5 μM). We treated the senescent cells with young ADSCs-sEVs (5 μg/mL), and we measured β-galactosidase (SABG) as a senescence marker and Annexin V as an apoptosis marker (Figure 3i). We found that cells that received ADSCs-sEVs showed a decrease in SABG, as well as a decrease in the percentage of apoptotic cells (Figure 3j-k).

Overall, these findings prove that treatment with ADSCs-sEVs can alleviate molecular and cellular traits of aging in kidney and muscle of aged mice, as well as a decrease in senescence and apoptosis in an *in vitro* model.

### Predicted epigenetic age is lower in tissues of old mice treated with ADSCs-sEVs

We used multiple mouse clocks trained on different tissues to estimate the age of our treated and untreated animals. We grouped the clocks into two groups. The general clocks were trained across a wide spectrum of animals. The interventions group of clocks were trained on animals with age affecting interventions. We used mixed effects modelling to discern a trend across multiple clocks. In kidney tissue samples, age estimates after the ADSCs-sEVs treatment were significantly lower for both general and interventions clocks (p = 0.006 and p = 0.0002, respectively). Similarly, in liver tissue samples, both models showed a reduction in estimated age with the interventions group passing the statistical significance threshold (p = 0.06 and p = 0.001 for general and interventions clocks, respectively). Muscle and spleen tissues did not show a significant difference in age prediction between treated and control animal groups (Figure 4a).

**Figure 4:**
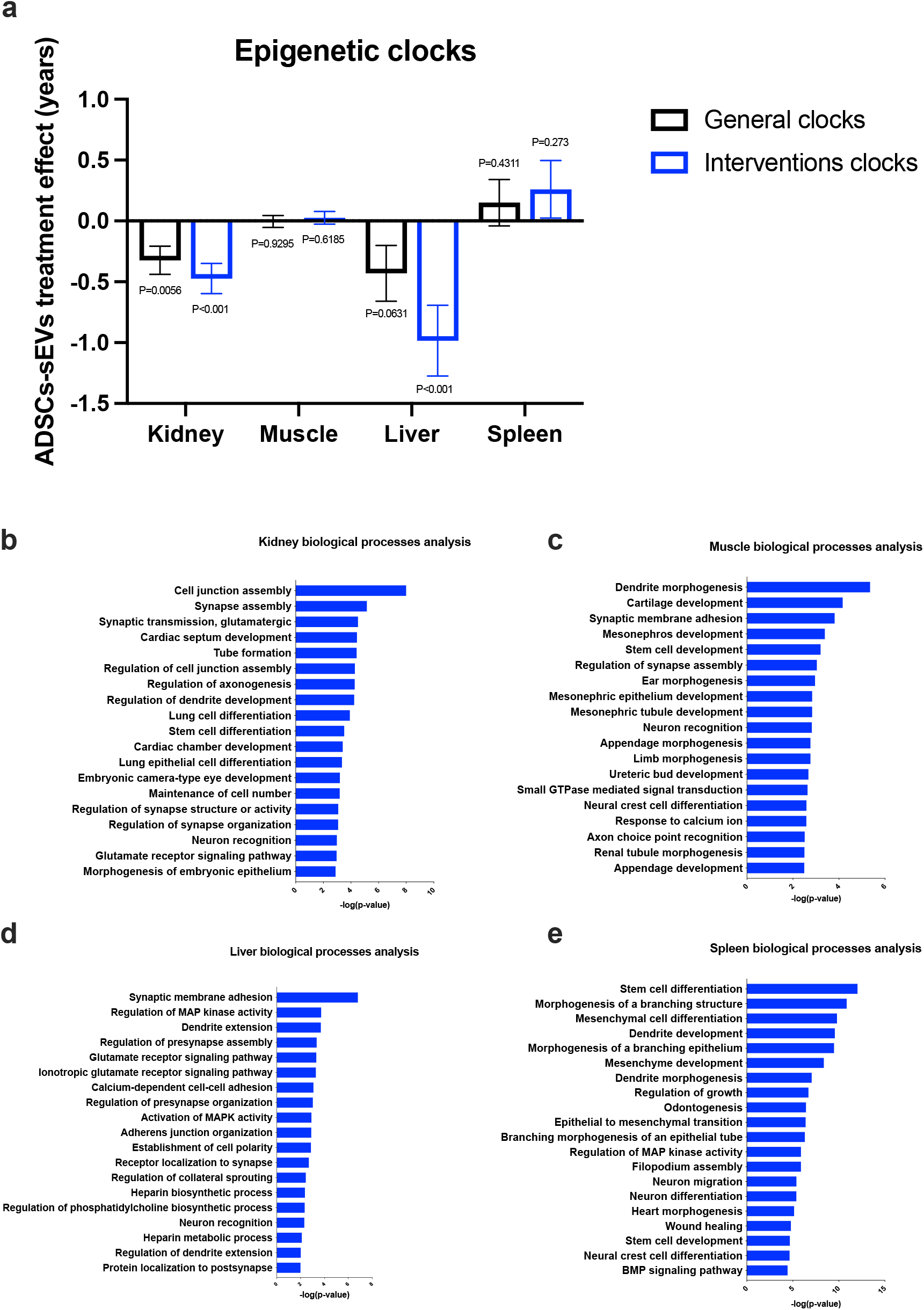
ADSCs-sEVs decrease the epigenetic age of several tissues in old mice. a: ADSCs-sEVs treatment effect on epigenetic age estimation mixed-effects modeling across several tissues. b-e: Biological processes enrichment analysis of differentially methylated cytosines in ADSCs-sEVs treated tissues. PBS n=6, ADSCs-sEVs n=6. All data are shown as mean±SD.

Linear regression analysis revealed abundant differentially modified loci in all tissues (n = 5.546, 36.163, 7.247, 11.092 in spleen, liver, kidney, muscle, respectively; robust linear regression, FDR q < 0.05). We assigned each locus the nearest gene name and performed gene set enrichment analysis using g:profiler^39^. Although affected pathways were different across tissues, we observed developmental, neuronal, and synaptic pathways similarly affected in all tissues (Figure 4b-e).

### ADSCs-sEVs induce a change in the metabolome of old mice to a youth-like pattern

We quantified 72 metabolites in blood plasma samples from 7 PBS-treated and 6 ADSCs-sEVs-treated old mice from 4 control young mice. Fourteen of these metabolites show statistically significant differences between ADSCs-sEVs-treated and untreated old animals (Supplementary Table 3 and Figure 5a). Most of the significant metabolites were amino acids including essential (isoleucine, tryptophan, threonine, and valine) and non-essential (aspartate, arginine, tyrosine, glycine, and proline). Some of these amino acids have been previously associated with metabolic health like the branched-chain amino acids isoleucine and valine^40^ as well as tryptophan^41^. Lactate is formed by the anaerobic glycolysis in most mammalian tissues. Together with acetate, it is also a product of fermentation of fructans and other prebiotics by gut microbiota^42^. They are further processed into butyrate and other short-chain fatty acids by some strains of *bifidobacteria* and *lactobacilli*, which suggest some impact of ADSCs-sEVs treatment in gut microbiota ecosystems. The heatmap of these metabolites shows a clustering trend among groups and some resemblance between ADSCs-sEVs-treated old mice and young mice (Figure 5b), i.e., red dots (old-sEVs mice) differ from black ones (old-PBS mice) but coincide with blue dots (young mice).

**Figure 5:**
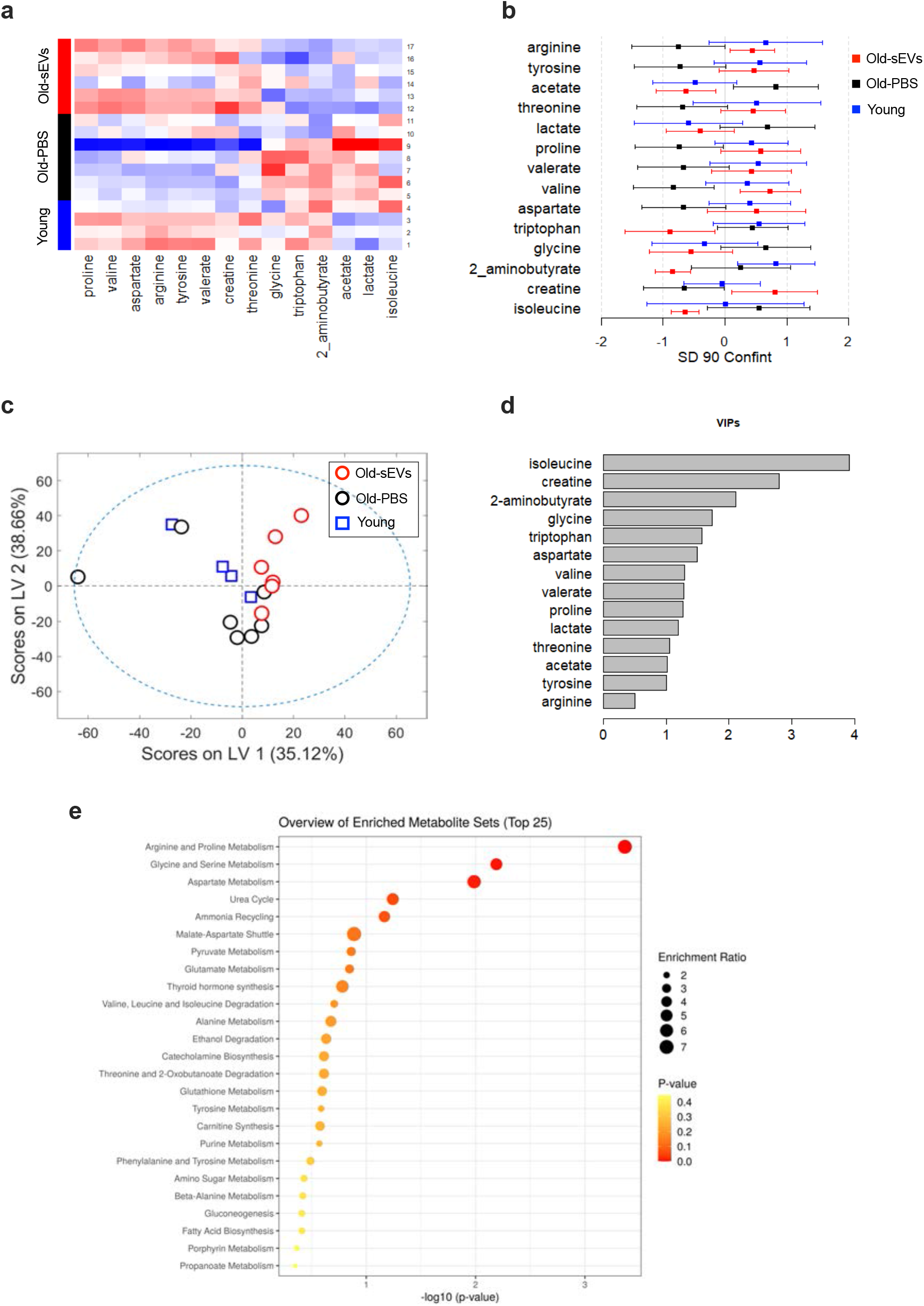
The metabolome of ADSCs-sEVs treated mice resembles that of young animals. a: Heatmap of metabolites z-scores with statistically significant differences between old-PBS (black bar samples) and old-sEVs (red bar samples) mice compared with young untreated mice (blue bar samples). b: Mean and 90% confidence intervals for z-scores in standard deviation units for the same metabolites that panel A in old-PBS (black), old-sEVs (red), and young untreated (blue) mice. c: Scores plot for a metabolome PLS-DA model (72 metabolites) for discrimination between young (blue squares) and old-PBS (black circles) mice. Old mice treated with sEVs (red circles) are projected in the scores plot. d: Variable importance in the projection (VIP) scores in the metabolome PLS-DA model of panel C for those metabolites significantly associated with treatment. e: Enrichment ratio (size of the circle) and p-values (colour of the circle) for the 25 most enriched metabolite sets from an MSE analysis on the SMPDB and our 14 significant metabolites. The n used for all metabolomic analysis was young mice n=4, old-PBS mice n=6 and old-sEVs mice n=6.

To further explore this resemblance, we built a PLS-DA model with all the metabolites for discriminating old and young untreated animals and projected into this model the samples from ADSCs-sEVs -treated animals. The scores plot of all the samples in this model shows the same resemblance between old treated and young animals along LV2 dimension suggesting that ADSCs-sEVs treatment partially ameliorates the impact of aging on the metabolome (Figure 5c). Thirteen out of the fourteen metabolites significantly associated with treatment also show a PLS-DA VIP score higher than 1, which confirms interactions between age-related metabolomic changes and ADSCs-sEVs treatment (Figure 5d).

Finally, we performed an MSE analysis concerning the Small Molecule Pathway Database (SMPDB) for mammal’s metabolism to gain biological insight into the metabolome changes induced by the ADSCs-sEVs treatment. The only three metabolite clusters enriched in our data with statistical significance were the Arginine and Proline metabolism cluster, the Glycine and Serine Metabolism cluster, and the Aspartate Metabolism cluster with 5, 4, and 3 hits, respectively (Figure 5e). These three clusters, especially the Arginine and Proline clusters, have been previously associated with cardiometabolic health in humans^43^. Besides the well-known role of arginine as urea or nitric oxide precursor, arginine is also a precursor of proline and relevant for collagen biosynthesis^44^. Proline and glycine, the two most abundant amino acids in collagen accounting for 23% of the molecule, are also represented in the first three clusters of our MSE analysis suggesting some involvement of collagen metabolism in the potential beneficial effects of ADSCs-sEVs treatment.

### Analysis of sEVs miRNA content reveals that upregulated miRNAs in young ADSCs-sEVs govern tissue development and regeneration pathways

Motivated by these results, we attempted to investigate candidate factors present in ADSCs-sEVs that could be responsible for the beneficial effects in old mice. miRNAs are an important fraction of several types of EVs content and can regulate transcription in several tissues ^45^. Here we used miRNA profiling (Affymetrix GeneChip miRNA 4.0 Array) to study miRNAs contained in sEVs. We obtained sEVs from three different conditions: supernatant of young and old ADSCs, and plasma from aged mice. In the analysis, miRNAs with a 2-fold increase or decrease in expression and a p-value <0,01 were defined as significantly different. Principal component analysis showed three differentiated groups of samples (Figure 6a). Analysis of 3196 miRNAs in sEVs from young ADSCs cultures showed several differentially expressed miRNAs when compared to sEVs from old ADSCs cultures (9 up-regulated, 1 down-regulated) and from plasma of aged mice (25 up-regulated, 4 down-regulated) (Figure 6b-d). In all comparisons, sEVs from young ADSCs showed an increased miRNA expression profile, indicating that probably there is a decreased regulation of miRNA biosynthesis with aging^46, 47^. These miRNAs are shown in Supplementary tables 4, 5, and 6.

**Figure 6:**
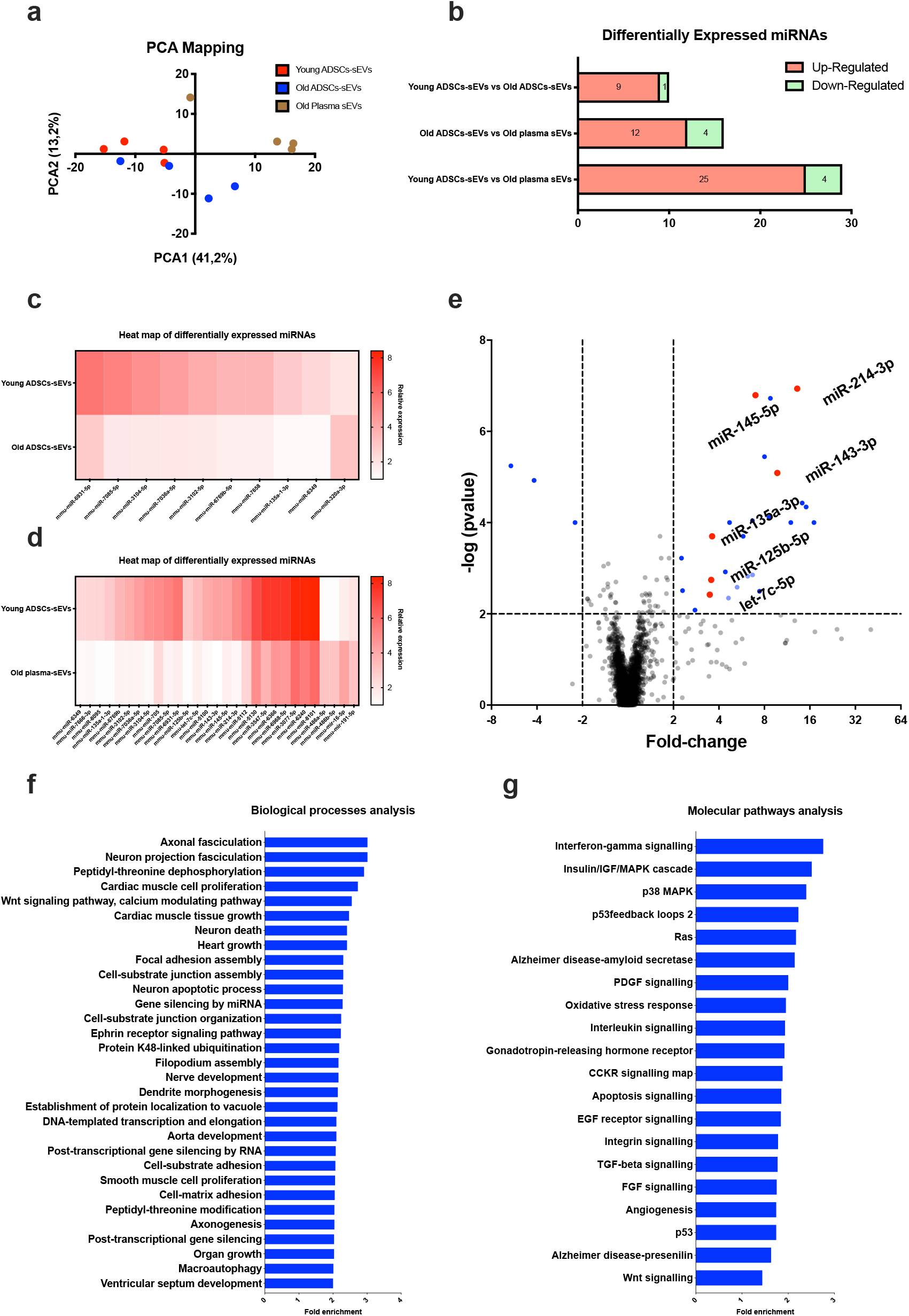
Analysis of sEVs miRNA content reveals that tissue development and regeneration are major targets of miRNAs-enriched in young ADSCs-sEVs. a: Principal component analysis of miRNA profiling in sEVs from young ADSCs, old ADSCs, and plasma of aged mice. Old plasma n=4, old ADSCs-sEVs n=4 and young ADSCs-sEVs n=4. b: Summary of differentially expressed miRNAs in the three conditions. c: Heat map of differentially expressed miRNAs in sEVs from young and old ADSCs. d: Heat map of differentially expressed miRNAs in sEVs from young ADSCs and plasma of aged mice e: Volcano plot showing the different miRNA expression between plasma of aged mice and young ADSCs-sEVs. f-g: Pathways and biological processes regulated by the target gene set of the six selected miRNAs. Images show the over-representation test run in PANTHER of the predicted targets of the miRNAs as defined by miRDIP (5379 genes).

As we injected the young ADSCs-sEVs in the blood of old mice, we reasoned that miRNAs upregulated in the young ADSCs-sEVs and with low levels in plasmatic sEVs from aged mice maybe, at least in part, responsible for the beneficial effects of ADSCs-sEVs. Thereafter, we explored the predicted mRNA targets for the upregulated 25 miRNAs and their role in biological processes and molecular pathways. However, as the number of predicted targets was too high, we outlined 6 miRNAs as plausible biologically relevant, these miRNAs (highlighted in red in Figure 6e) were the ones that shared their nucleotide sequence among several species, including humans, as many features of the aging process are highly conserved across species^48^.

Altogether, these 6 miRNAs were predicted to target 5.379 genes with very high stringency (top 1% in confidence, present in at least five databases as curated by miRDIP). The target gene dataset was then introduced in PANTHER to study over-represented biological processes and pathways ^48^. A great number of genes targeted by our miRNAs were involved in cell proliferation, tissue development, growth, and regeneration (Figure 6f). Regarding molecular pathways, these miRNAs regulated important pathways related to tissue regeneration, fibrosis, and inflammation, such as angiogenesis, Wnt signalling, PDGF, FGF, TGF-β, interleukin, and Interferon-γ signalling. Most importantly, they targeted critical pathways in the aging process, including apoptosis, p53 signalling, p38 MAPK, oxidative stress response, and insulin/IGF pathway (Figure 6g). In conclusion, predicted targets of the selected miRNAs upregulated in young ADSCs-sEVs compared with plasma sEVs of old mice, are involved in several relevant processes and pathways affected during aging.

## Discussion

Aging is the largest risk factor for most diseases that affect people at late stages of their lives. Our understanding of the aging process has grown greatly in the last decades, providing us with several biochemical changes that are considered as drivers of this process^2^. An altered intercellular communication is one of the ultimate culprits of tissue and organismal aging and several players of intercellular communication have been proposed as possible anti-aging factors^6, 7, 49^. sEVs could be one important fraction of the factors present in young environments that can have a beneficial effect on aged individuals. sEVs from MSCs have a regenerative capacity themselves, they can help tissue to regenerate after damage, and are potential cell-free therapeutic agents^50^.

Nicotinamide phosphoribosyl transferase (eNAMPT) was shown to be present within plasmatic EVs and these EVs were able to ameliorate several parameters affected by aging and increased lifespan of old mice^25^. Similarly, sEVs have been described as carriers of glutathione-S-transferase activity, which can ameliorate senescence and oxidative stress-related damage in old mice^23^. More recently, sEVs from MSCs were able to extend lifespan in a transgenic mouse model of accelerated aging and mitigated several senescence-associated markers *in vivo* and *vitro*^51^.

Given this background, the effect of sEVs from young MSCs on the function of tissues affected by aging in physiologically aged mice remained to be elucidated. In the present work, we have shown that young-derived ADSCs-sEVs have great potential as anti-aging factors, as old mice injected with these sEVs showed an improvement in several functions affected by aging, such as physical condition, fur regeneration, and renal function, as well as a reduction of frailty. Very interestingly, the effect of ADSCs-sEVs seems to be finite, as we observed in a more long-term experiment that the effect on the functionality of old mice was lost two months after treatment with ADSC-sEVs. Moreover, sEVs induced structural changes in tissues from old mice, exhibiting a potent pro-regenerative effect in these tissues, which is in accordance with previous results that proposed sEVs as a regenerative tool^23^. We observed a reduction of senescence in tissues and *in vitro* when sEVs were introduced, however, the mechanism of action remains unclear, as we did not find senolytic activity. They may probably act as senomorphics, molecules that suppress the senescent phenotype, as has been described recently^51^. Indeed, we found a reduced level of IL-6, one of the main SASP factors, in the tissues from ADSCs-sEVs treated mice. All these effects found in tissues of ADSCs-sEVs treated mice were accompanied by a decrease in the epigenetic age estimation from epigenetic clocks, which are recognized as a feasible estimation of biological age^52^. Some of the most studied interventions in aging, such as caloric restriction or rapamycin treatment, have demonstrated a strong impact on predicted epigenetic age ^53,54,55^. To our knowledge, this study is the first to show that intervention with sEVs can decrease epigenetic age in an old organism.

sEVs induced effects not only in individual tissues but also in the whole metabolome of old mice, which turned to a youth-like pattern, indicating that the effects observed are probably part of a pleiotropic effect on the organism. Now as it is time to continue investigating the molecules that are included in sEVs, we have explored miRNAs contained in young ADSCs-sEVs and found that they are involved in several processes and pathways affected by aging, thus proposing miRNAs as possible mediators of the effects shown in mice, miR-214-3p seems to play a key role in senescence^56, 57^. More studies are needed to identify factors derived from stem cells that can assist tissue function and regeneration, as they could have an enormous impact on age-related pathologies, such as frailty or renal failure.

## Supporting information

Mice video

## Acknowledgments

We appreciate the contribution of the Flow Cytometry Service, Multigenic Analysis Unit, Animal Facility, and Metabolomic and Molecular Image Laboratory of the Central Unit in Medical Research, as well as the Microscopy section of the Central Service of Support to the Experimental Research, of the University of Valencia. We also want to thank the help provided by the Electronic Microscopy service of the Príncipe Felipe Research Centre in Valencia. We also thank Mrs. Marilyn Noyes for her kind help in reviewing the manuscript. This work was supported by the following grants: Grant PID2020-113839RB-I00 funded by MCIN/AEI/ 10.13039/501100011033, PCIN-2017-117 of the Ministry of Economy and Competitiveness, and the EU Joint Programming Initiative ‘A Healthy Diet for a Healthy Life’ (JPI HDHL INTIMIC-085) to CB and CB16/10/00435 (CIBERFES) from Instituto de Salud Carlos III, (PID2019-110906RB-I00/ AEI / 10.13039/501100011033) and RED2018-102576-T from the Spanish Ministry of Innovation and Science, PROMETEO/2019/097 from “Consellería, de Educació de la Generalitat Valenciana” and EU Funded H2020-DIABFRAIL-LATAM (Ref: 825546), European Joint Programming Initiative “A Healthy Diet for a Healthy Life” (JPI HDHL) and of the ERA-NET Cofund ERA-HDHL (GA N° 696295 of the EU Horizon 2020 Research and Innovation Programme) and Fundación Ramón Areces y Fundación Soria Melguizo. to J.V., and PID2019-108973RB-C22 to D.M. Part of the equipment employed in this work has been funded by Generalitat Valenciana and co-financed with ERDF funds (OP ERDF of Comunitat Valenciana 2014-2020).

## Author contributions

J.S-R, J.V, and C.B conceived and designed the study. J.S-R conducted most experiments, C.M-B, A.R-D, N.R-G, and M.I assisted the MDA determination assay. A.D and A.G-C performed telomere assays. D.M performed the metabolomic assay. R.T.B. performed epigenetic clocks. J.S-R, A.D and A.G-C, D.M and J.G. analyzed the data. J.S-R, D.M, J.G., and C.B wrote the manuscript, M.A.B, S.H., J.V, and C.B revised the manuscript.

## Materials and methods

### Animal model

This study was performed in strict accordance with all applicable federal and institutional policies. The protocol was approved by the University of Valencia Animal Ethics Committee. All the mice used in this study were of a C57BL/6J background. Aged mice used were treated at 22-24 months of age. Both sexes were used throughout the study. Where feasible, littermates of the same sex were used. These were randomly assigned to experimental groups. All experiments were addressed blindly. After mice were sacrificed, plasma and organs were obtained and stored at −80°C. For all the experiments conducted in kidney and gastrocnemius, we used organs obtained from mice 30 days after treatment with sEVs/PBS.

### Stem cell culture

Mice at 3-6 months of age were used to obtain mesenchymal stem cells (MSCs) from both inguinal fat pads with the previously described protocol ^58^. All cells were used at passage 2-3 for the experiments. The culture media used for the expansion and maintenance of the cells was DMEM high glucose with 10% of foetal bovine serum (FBS) and 1% of Penicillin/Streptomycin (P/S). Cells were maintained at 37°C, 5% CO2, and 3% O2 (Whitley H35 HEPA Hypoxystation).

### ADSCs characterization

For the characterization of ADSCs we analyzed the presence of typical MSCs markers (CD29, CD44, CD90, CD105) by flow cytometry, CD45 and CD31 were used as negative controls. When cells reached 80% confluency, they were trypsinized and resuspended in FACS buffer (10% FBS, 1% sodium azide in PBS). 100,000 cells were used for each condition, they were stained with 1 µg/µL of antibody and incubated for 30 minutes at 4°C. All antibodies were acquired from Biolegend: CD29 PE/Cy7, 10222; CD31 PE, 102507; CD44 Alexa 488, 103015; CD45 PE, 103106; CD90 FITC, 140303; CD105 APC, 120413).

ADSCs isolated from inguinal fat pads of young mice showed a strong expression of MSCs markers, cells were positive (>80%) for CD29, CD44, CD9,0, and CD105 and negative for hematopoietic (CD45) and endothelial (CD31) markers (Supplementary figure 1).

### ADSCs derived sEVs isolation and characterization

ADSCs at passage 2-3 were expanded until they reached 80% confluency, at that moment, media was changed to DMEM high glucose with 2% Exosome-depleted FBS (Gibco) and 1% P/S. After 48h, conditioned media was collected, and sEVs were isolated by differential ultracentrifugation. Media was centrifuged at 2,000g for 10 minutes and then at 20,000g for 30 minutes to remove whole cells, cell debris, and bigger EVs. The supernatant was then ultracentrifuged at 100,000g for 70 minutes. Pelleted vesicles were suspended in PBS, ultracentrifuged again at 100,000g for 70 minutes for washing, resuspended in P, BS, and prepared for treatment, electron microscopy, or flow cytometry analysis.

For the characterization of sEVs isolated from the conditioned media, we checked for the expression of CD63, a ubiquitous marker in the membrane of sEVs^59^. Using flow cytometry, we were able to confirm the presence of this marker in the samples (Supplementary figure 1). For flow cytometry analysis pellets obtained from 10 mL of media were resuspended in 100 µL PBS. sEVs were stained with 4µg/mL of APC anti-CD63 (Biolegend, 143905) for 30 minutes at 4°C in dark. After incubation, positive events were read by FACS-Verse flow cytometry. One unstained sample of sEVs and one sample with the antibody at 4 µg/mL in PBS were used as negative controls. Calibration of the cytometer was assessed using fluorescent particles of standardized size (Nano Fluorescent Particle Size Standard Kit, Spherotech).

Secondly, we used transmission electron microscopy to measure the size and morphology of the isolated vesicles. The presence of round-shaped vesicles that ranged from 50-200 nm in diameter was confirmed, confirming that the isolate was largely enriched in sEVs (Supplementary figure 1). The isolated sEVs were fixed in 2 % Paraformaldehyde - 0.1 M phosphate-buffered saline for 30 min. Glow discharge technique (30 sec, 7,2V, using a Bal-Tec MED 020 Coating System) was applied over carbon-coated copper grids, and immediately, these grids were placed on top of sample drops for 15 min. Then, the grids with adherent sEVs were washed in a 0.1, M PB drop and additional fixation in 1 % glutaraldehyde was performed for 5 min. After washing properly in distilled water, the grids were contrasted with 1 % uranyl acetate and embedded in methylcellulose. Excess fluid was removed and allowed to dry before examination with a transmission electron microscope FEI Tecnai G2 Spirit (ThermoFisher Scientific company, Oregon, USA). All images were acquired using a digital camera Morada (EMSIS GmbH, Münster, Germany).

Ultimately, we assessed the presence of CD63 on the membrane of the sEVs through immunogold staining (Supplementary figure 1). In brief, 8 μl of isolated sEVs were fixed in 2 % Paraformaldehyde - 0.1 M phosphate-buffered saline for 30 min, and carbon-coated nickel grids were placed on top of these sEVs drops for 15 min. Then, the grids with adherent sEVs were washed in 0.1 M PBS and blocked in 0.1 M glycine and 0.3 % BSA for 10 min. The grids were incubated with anti-CD63 (MBL, D263-3) primary antibody (1:100 dilution) for 1 hour. After a new blocking step for 10 min, the grids were incubated in Gold 6nm conjugated Goat anti-rat (Abcam, ab105300) secondary antibody (1:1000 dilution) for 1 h. Finally, after washing, a standard negative staining procedure was made and observed under a Transmission Electron Microscope as previously described.

### Treatment preparation

sEVs isolated from the conditioned media were kept at 4°C and used for treatment within 24 hours. To determine the dose of sEVs, we performed protein quantification with the Lowry method of the sEVs samples. Each dose consisted of either 20µg of sEVs’ protein or PBS as a control in a total volume of 100µL.

### Physical condition tests

The Grip Strength Meter (Panlab, Harvard Apparatus) was employed in assessing neuromuscular function by sensing the peak amount of force that the mice applied in grasping specially designed pull bar assemblies. Peak force was automatically registered in grams-force by the apparatus. Data were recorded, and 2 additional trials were immediately given. Maximum force was normalized to the weight of the mouse. Animals were submitted to a graded intensity treadmill test (Treadmill Control LE 8710 Panlab, Harvard Apparatus) to determine their endurance (running time) and running speed along with the study. After a warmup period, the treadmill band velocity was increased until the animals were unable to run further. The initial bout of 4 minutes at 10 cm/s was followed by consecutive 4 cm/s increments every 2 minutes. Exhaustion was reached when a mouse remained on the shock grid for 5 seconds rather than running. Motor coordination was assessed with a Rota Rod (Panlab, Harvard Apparatus #76-0772), consisting of a 3cm wide wheel that rotates with an increasing speed. Time to fall was recorded for each mouse, with a total of 3 trials. To assess frailty quantitatively, we used a score based on the clinical phenotype of frailty developed in our group^26^, mice with 3 or more criteria were classified as frail. Criteria used: Loss of 5% of body weight, time to fall in Rota Rod under percentile 20 (p20), time to exhaustion in a treadmill under p20, maximum speed reached in a treadmill under p20, and normalized grip strength under p20.

### Hair re-growth assay

On day 1, dorsal hair was removed by plucking from a square of approximately 1 cm x 1 cm. Hair re-growth was scored two weeks later, based on digital photographs and a semi-quantitative assessment, using an arbitrary scale from one to four (where four represents complete hair regeneration). Scoring was performed blindly by two independent investigators.

### Plasmatic urea values measurement

100 µL of whole blood was obtained from the saphenous vein of each mouse just before treatment, after that, whole blood was obtained on days 14, 30, and 60. Whole blood samples were collected in a microvette with Ethylenediaminetetraacetic acid (EDTA) for plasma separation and spun for 15 minutes at 1,500g. The clear supernatant was transferred into regular 1.5ml tubes, snap-frozen in liquid N2, and stored at −80°C. [Urea] was measured using a QuantiChrom Urea Assay Kit (Gentaur). The samples were incubated in a 200µl reaction mix for 20 min at room temperature before absorbance was measured at 520nm.

### Histological analysis

Kidneys and gastrocnemius were fixed in 4% PFA for 48h and then cryoprotected with 30% sucrose in PBS, 10µm slices were obtained in a cryostat for histological staining, and immunofluorescence. Slices were stained with Haematoxylin (Sigma, MHS32) and Eosin (Sigma, E4009), or Sirius Red (Sigma, 365548), mounted, and sealed for further morphometric analysis. Images were obtained using an optical microscope (Leica), three images from different areas of each slice were obtained. All levels were adjusted equally, and the ratios were not altered, morphometric analysis of kidney and muscle sections was performed with ImageJ.

### Immunofluorescence

10µm tissue slices were mounted in slides and heated for 30 minutes in Tris-EDTA buffer pH 10 for antigen retrieval and permeabilized with 1% TritonX-100 in PBS (Ki-67) or 1% Tween 20 in PBS (LMNB1), depending on the antibody. Sections were blocked with 10% normal goat serum (Invitrogen) in PBS 0,05% Tween 20 (PBS-t) and incubated with primary antibody overnight at 4°C. After incubation with primary antibody, sections were washed 3 times with PBS-t and incubated with secondary antibody for 2 hours at RT, washes were repeated, and tissues were counterstained with Hoechst (Invitrogen) for 30 minutes at RT. Coverslips were mounted with an aqueous mounting medium and sealed with nail polish. Images were acquired with Olympus FV1000 confocal laser scanning biological microscope. Images were processed in ImageJ all levels were adjusted equally and the ratios were not altered. Ki67 staining antibodies: anti-Ki67 (ThermoFisher, PA5-19462, 1:1000 dilution) and AlexaFluor 647 anti-rabbit (Abcam, ab150079, 1:1000 dilution). LMNB1 staining antibodies: anti-LMNB1 (Proteintech, 12987-1-AP, 1:50 dilution) and AlexaFluor 488 anti-rabbit (Abcam, ab150079, 1:2000 dilution).

### Lipid peroxidation measured using high-performance liquid chromatography

Tissues were lysed with a KPi-EDTA buffer (KPi 50mM, EDTA 1mM pH 7.4), lipid peroxidation was determined as malondialdehyde (MDA) levels, which were detected using high-performance liquid chromatography (HPLC) as an MDA-thiobarbituric acid (TBA) adduct following a method described previously. This method is based on the hydrolysis of lipoperoxides and subsequent formation of an adduct between TBA and MDA (TBA-MDA2). This adduct was detected using HPLC in the reverse phase and quantified at 532 nm. The chromatographic technique was performed under isocratic conditions, the mobile phase being a mixture of monopotassium phosphate 50 mM (pH 6.8) and acetonitrile (70:30). The level of MDA in each sample was divided by the concentration of protein determined by the Lowry method.

### Immunoblotting

Tissues and cells were lysed in Tris/SDS/Glycerol buffer, proteins were separated on SDS polyacrylamide gels and transferred onto nitrocellulose membranes. The membranes were blocked with 3% BSA in Tris-buffered saline, 0,05% Tween 20 (TBS-t) for 60 min at RT and incubated overnight at 4°C with anti-IL6 antibody (Abcam, ab229381, 1:1000 dilution). Following 3 washes (10 min) with TBS-t, membranes were incubated with secondary antibody (Cell Signalling, 70745, 1:2000 dilution) for 60 min at RT. Following 3 washes with TBS-t membranes were developed with Luminol (Sigma) in the ImageQuant LAS4000 system. Images were processed in ImageJ. To detect total protein carbonylation by immunoblot we used the OxyBlot Protein Oxidation Detection kit (Merck) following the manufacturer’s instructions. Ponceau staining of the membranes was used as the loading control.

### Telomere length and telomere damage assays

Telomere length and the number of telomere dysfunction-induced foci (TIF) were calculated by means of the IF-FISH technique. Briefly, frozen slides of the kidney and muscle were permeabilized with 0.5% Triton X-100 and washed in PBS. After 1 hour of blocking with 5%BSA-PBS, slides were incubated with 53BP1 antibody (Novus Biologicals, NB100-304) at 4C, o/n. After several washes in PBS-0.1% Tween20, slides were incubated in donkey anti-rabbit Alexa Fluor 488 antibody (Invitrogen, A21206) for an hour and washed again in PBS. Tissues were then fixed in 4% formaldehyde for 20 min, washed several times in PBS, and dehydrated in a 70% - 90% - 100% ethanol series (5 min each). Slides were left to air dry and 30ul of telomere probe mix -10mM TrisCl pH7, 25mM MgCl2, 9mM Citric Acid, 82 mM Na2HPO4, 50% Deionized Formamide (Ambion AM9342), 0.25% Blocking Reagent (Roche 11096176001), and 0.5μg/ml Telomeric PNA-Cy3 probe (Panagene)-were added to each slide. A coverslip was added, and slides were incubated for 3 min at 85°C and further 2h at RT in a wet chamber in the dark. Slides were washed 2×15 min in 10 mM TrisCl pH7, 0.1% BSA in 50% formamide under vigorous shaking, then 3×5min in TBS 0.08% Tween20 and rinsed in PBS. Slides were mounted in Vectashield with DAPI (VectorTM H-1200-10). Confocal images were acquired as stacks using a Leica SP5-MP confocal microscope and maximum projections were done with the LAS-AF Software. Telomere signal intensity was quantified using Definiens Software and sites of colocalization of telomeric PNA-Cy3 probe and 53BP1 fluorescent signals were counted per cell. Image analysis was performed blindly.

### Epigenetic age estimation using epigenetic clocks

DNA from different tissues was obtained using Qiagen DNeasy Blood & Tissue Kit, following the manufacturer’s instructions. Raw files were processed using R package sesame version 1.3.0. The beta values were obtained using the default sesame procedure with nondetection.mask and quality.mask set to FALSE. Differentially methylated cytosines were detected using robust linear regression implemented in R package limma separately for each tissue. The treatment effect was estimated for each CpG by adjusting for potential confounding from age, sex, and sentrix array row. Obtained p values were corrected for multiple testing using the FDR procedure and those with q < 0.05 were deemed significant. Pathway enrichment analysis was performed by annotating the loci to the nearest gene and ranking them by signed log p-value. The ranked list was passed to g:profiler for enrichment analysis using R package gprofiler2^39^.

Multiple mouse age predictors (epigenetic clocks) were used to estimate the age of the animals. The mouse clocks were trained/developed in a separate data set ^60^. The age predictors were grouped into general and intervention clocks. In the general group, the clocks were trained on the following mouse tissues across a wide spectrum of animals: blood, liver, brain, cerebellum, cortex, fibroblast, heart, kidney, muscle, skin, striatum, tail, and a pan tissue clock spanning all different tissues. In the interventions group, the clocks were trained across the same tissues and animals, but the loci used for training were those known to change methylation levels upon known age affecting interventions. The effect of ADSCs-sEVs treatment was estimated by fitting a mixed effect model for each tissue and group of clocks where animal age, sex, and treatment were fixed variables and clock name was a random variable. Age predicted by the clock was the response variable for the model. The statistical significance of each fixed variable was evaluated using Satterthwaite’s method implemented in R package lmerTest. Array normalization, array manifest file, array annotation files, epigenetic clock coefficients were obtained from Steve Horvath’s lab.

### Plasma metabolites by Proton Nuclear Magnetic Resonance (1H-NMR) Spectroscopy

For each sample, a mixture of 20 uL of blood plasma and 2 uL of phosphate buffer with trimethylsilyl propanoic acid (TSP) and deuterated water was transferred into a 1 mm high-quality NMR individual tube. Proton NMR spectra for all samples were recorded in a Bruker Avance DRX 600 spectrometer, equipped with a triple resonance 1H/13C/31P probe. The nominal temperature of the sample was kept at 310 K. A single pulse presaturation experiment was acquired in all samples. The number of transients was 256 collected into 65 k data points for all experiments. Spectral chemical shift referencing on the alanine CH3 doublet signal at 1.475 ppm was performed in all spectra. Spectra were processed using MestReNova 8.1 (Mestrelab Research S.L., Spain) and transferred to MATLAB (MathWorks, 2012) using in-house scripts for data analysis. The chemical shift region including resonances 0.50-4.70 ppm (the aliphatic region) and 5.20-10.00 ppm (the aromatic region) was investigated. Metabolite spin systems and resonances were identified by literature data and Chenomx resonances database (Chenomx NMR 7.6). Spectra were normalized to the total aliphatic spectral area, lipid excluded, to eliminate differences in metabolite total concentration. NMR peaks were integrated and quantified using semi-automated in-house MATLAB peak-fitting routines. Final metabolite levels were calculated in arbitrary units as peak area normalized to the total spectral area. Chemometrics analysis was performed with PLS ToolBox 8.0 (Eigenvector Inc) in MATLAB. To maximize the separation between samples and to identify discriminant patterns, partial least-squares discriminant analysis (PLS-DA) was applied. The permutation test was performed to check the overfitting of the PLS-DA models. The multivariate chemometric models were cross-validated with 10-fold Venetian blind cross-validation. In each run, 10% of the data were left out of the training and used to test the model. Spectral regions with high variable importance in projections (VIP) coefficients obtained during PLS-DA are more important in providing class separation during analysis, while those with very small VIP coefficients provide a little contribution to classification. A Metabolite Set Enrichment Analysis (MSEA) over metabolites with VIPs scores higher than 0.5 and p-values below 0.05 was performed with MetaboAnalyst and the Small Molecule Pathway Database (SMPDB). MSEA is conceptually like Gene Set Enrichment Analysis and uses a collection of predefined metabolites sets to rank the lists of metabolites obtained from metabolomics studies. By using this prior knowledge about metabolite sets, we could identify significant and coordinated changes in metabolic networks and obtain biological insight.

### Total RNA extraction from sEVs and small non-coding RNA expression profiling

sEVs obtained from different sources were stored in PBS at −80°C, for the isolation of RNA, we used the Total Exosome RNA Protein isolation kit (ThermoFisher), following the manufacturer’s instructions.

Small non-coding RNA expression profiling was performed using a GeneChip miRNA 4.0 Array (Affymetrix, Santa Clara, CA). Microarray experiments were conducted according to the manufacturer’s protocol. Briefly, 130 ng of total RNA was labeled with a FlashTag Biotin HSR RNA labeling kit from Genisphere. The labeling reaction was hybridized onto the miRNA array in an Affymetrix hybridization oven 645 at 48°C for 18 h. The arrays were stained using a Fluidics Station 450 with the fluidics script FS450_0002 (Affymetrix) and then scanned on a GeneChip Scanner 3,000 7G (Affymetrix, Santa Clara, CA), using the GeneChip Command Console© Software supplied by Affymetrix to perform gene expression analysis. MiRNA probe outliers were defined and further analyzed as per the manufacturer’s instructions (Affymetrix), and quality control, as well as data summarization and normalization, was carried out using the web-based miRNA QC Tool (www.affymetrix.com).

miRNA expression levels were analyzed with Transcriptome Analysis Suite (ThermoFisher). Data (.CEL files) were analyzed and statistically filtered using Transcriptome Analysis Suite (ThermoFisher), and input files were normalized with the multi-array average algorithm (RMA) for miRNAs. A one-way analysis of variance (ANOVA) was performed with the Transcriptome Analysis Suite (ThermoFisher) on all samples. Statistically significant small non-coding RNAs between the different groups and treatments studied were identified using an analysis of variance model with a P-value of 0.01 or less. The imported data were analyzed by principal components analysis (PCA) to determine the significant sources of variability in the data; PCA reduces the complexity of high-dimensional data and simplifies the task of identifying patterns and sources of variability in a large data set.

### miRNA pathway and biological processes analysis

From the upregulated miRNAs comparing sEVs from young ADSCs to old mice’s plasmatic sEVs, the 6 miRNAs conserved across species were analyzed using mirDIP to determine miRNA target genes with very high confidence (top 1% in confidence). A total of 5379 unique genes were further analyzed by PANTHER to identify enriched biological pathways and biological processes with an overrepresentation test, in PANTHER, Fisher’s exact test was used with False discovery rate correction. P-value<0,01 was considered statistically significant. For the biological process analysis, we considered only processes with 2-fold enrichment or more.

### *In vitro* model of senescence in muscle progenitor cells

For the *in vitro* experiments, we used the cell line C2C12 (ATCC CRL-1772), a mouse myoblast established line. The culture media used for the expansion and maintenance of the cells was DMEM high glucose with 10% of foetal bovine serum (FBS) and 1% of Penicillin/Streptomycin (P/S). Cells were maintained at 37°C, 5% CO2. Cells were seeded at a 10.000 cells/cm^2^ density, and 24 h after seeding, they were treated with 5µM Palbociclib for 96h. 72h after seeding, myoblasts were treated for 48 hours in EVs-depleted culture medium with 5µg/ml of ADSC-sEVs in PBS or PBS alone as a control. After that, cells were collected for further analysis by flow cytometry.

### Flow cytometry analysis of C2C12 cells

To determine cellular levels of SABG we used FluoReporter lacZ Flow Cytometry Kit (Invitrogen) following the manufacturer’s instructions. In brief, we harvested cells from the cell culture plate and diluted them to 1000 cells/µL in a staining medium, and introduced a 1:1 volume of fluorescein di-β-D-galactopyranoside (FDG) 2 mM working solution, cells were stained for exactly one minute at 37 °C. FDG loading was interrupted by adding 1.8mL of ice-cold staining medium containing 1.5μM propidium iodide. Fluorescence values were read by FACS–Verse flow cytometry (BDBiosciences, San Diego, CA, USA) until 10000 events were recorded.

Apoptosis was assessed using Annexin V FITC Apoptosis detection kit (Immunostep, Salamanca, Spain), following the manufacturer’s instructions. ADSCs were harvested and diluted to 1000 cells/µL in Annexin-binding buffer. Then, 100 µL aliquots of resuspended cells were placed into an appropriate flow cytometer tube and stained with 5 µL Annexin V-FITC and 5 µL propidium iodide for exactly 15 min at 37 °C in darkness. After incubation, 400 µL of 1 × Annexin-Binding Buffer were added. The values were read by FACS–Verse flow cytometry (BDBiosciences, San Diego, CA, USA) until 10000 events were recorded.

### Statistical analysis

Ratios comparing physical test and plasma values after treatment compared to baseline were determined and plotted as % over baseline. The baseline was defined as 0%. All groups were tested for the presence of outliers with the ROUT method (Q=2%). Saphiro-Wilk test was conducted in each comparison to test the normality of each group. Unpaired Student’s t-test or Mann-Whitney test were used to calculate the p-value for pairwise comparisons. For the in vitro experiments, ANOVA was used with Tukey’s multiple comparison as a post-hoc test. GraphPad Prism 9.0 software was used for the analysis.

## Supplementary information

**Supplementary figure 1:**
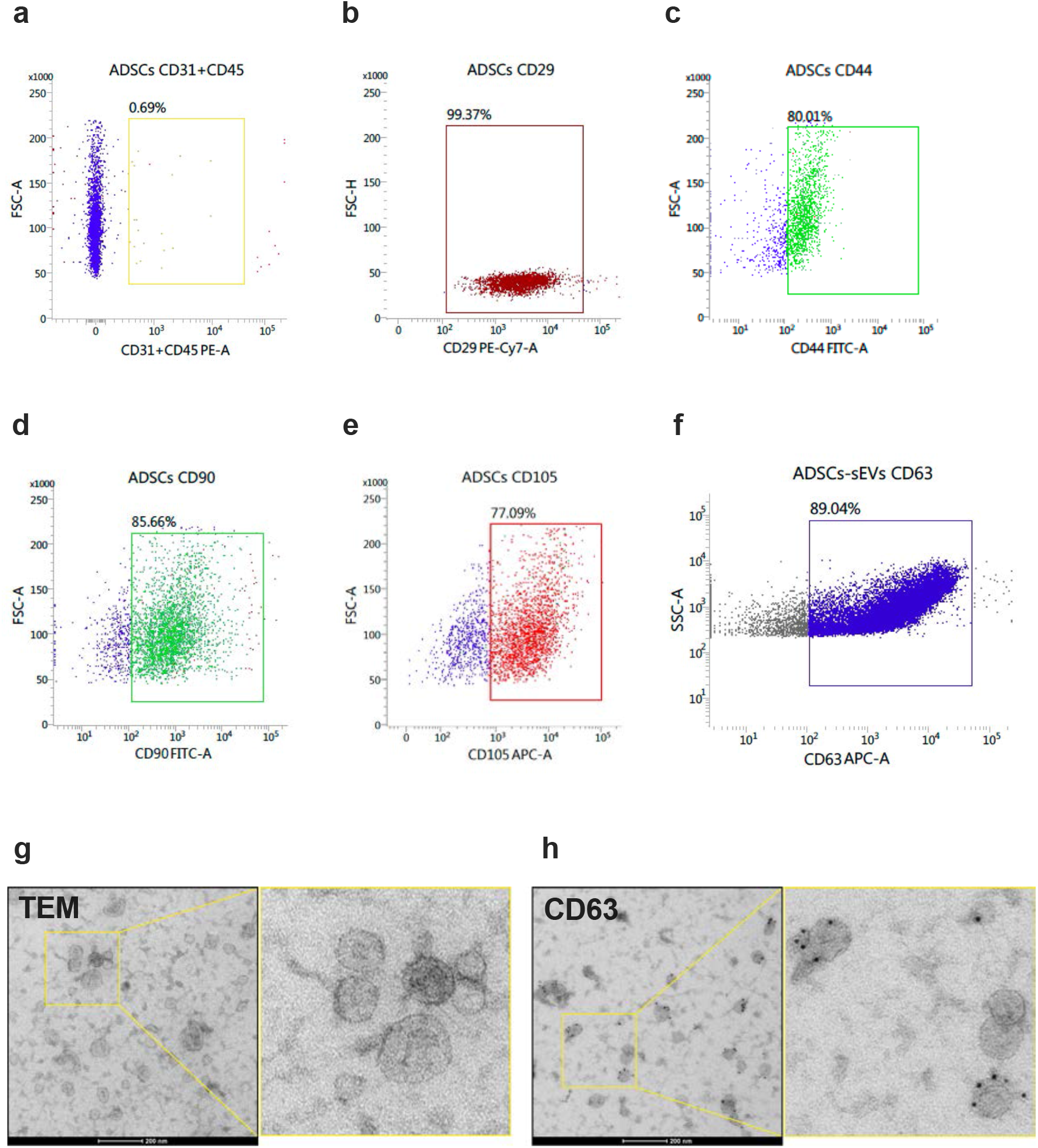
ADSCs release sEVs to the media. a-e: Quantification of mesenchymal stem cell markers in cells isolated from adipose tissue of young mice. f: Determination of CD63 presence in sEVs obtained from cell culture media of young ADSCs. g-h: Representative image of ADSCs-sEVs as observed by TEM and CD63 determination with immuno-gold labeling.

**Supplementary Table 1:** Data complementary to the study shown in Figure 1. Mice received two doses (on days 1 and 7 of the experiment) of sEVs released by ADSCs isolated from young mice (4-6 months). Results are shown in percentage of increase or decrease from baseline at day 60 of the experiment. Non-significant: ns.

| Test (day 60) | PBS | Young sEVs | PBS vs Young sEVs |
| --- | --- | --- | --- |
| Grip strength | +1,7% | +2,09% | ns |
| Motor coordination | -12% | +11% | ns |
| Fatigue resistance | -27,08% | -25,07% | ns |

**Supplementary Table 2:** Data of a pilot study with n=3 mice injected with two doses (on days 1 and 7 of the experiment) of sEVs released by ADSCs isolated from aged mice (24 months). Results are shown in percentage of increase or decrease from baseline at day 30 of the experiment. Non-significant: ns.

| Test (day 30) | PBS | Old sEVs | PBS vs Old sEVs |
| --- | --- | --- | --- |
| Grip strength | +4,23% | +6,61% | ns |
| Motor coordination | -2,89% | +1,99% | ns |
| Fatigue resistance | -16,36% | -10,03% | ns |

**Supplementary Table 3:**
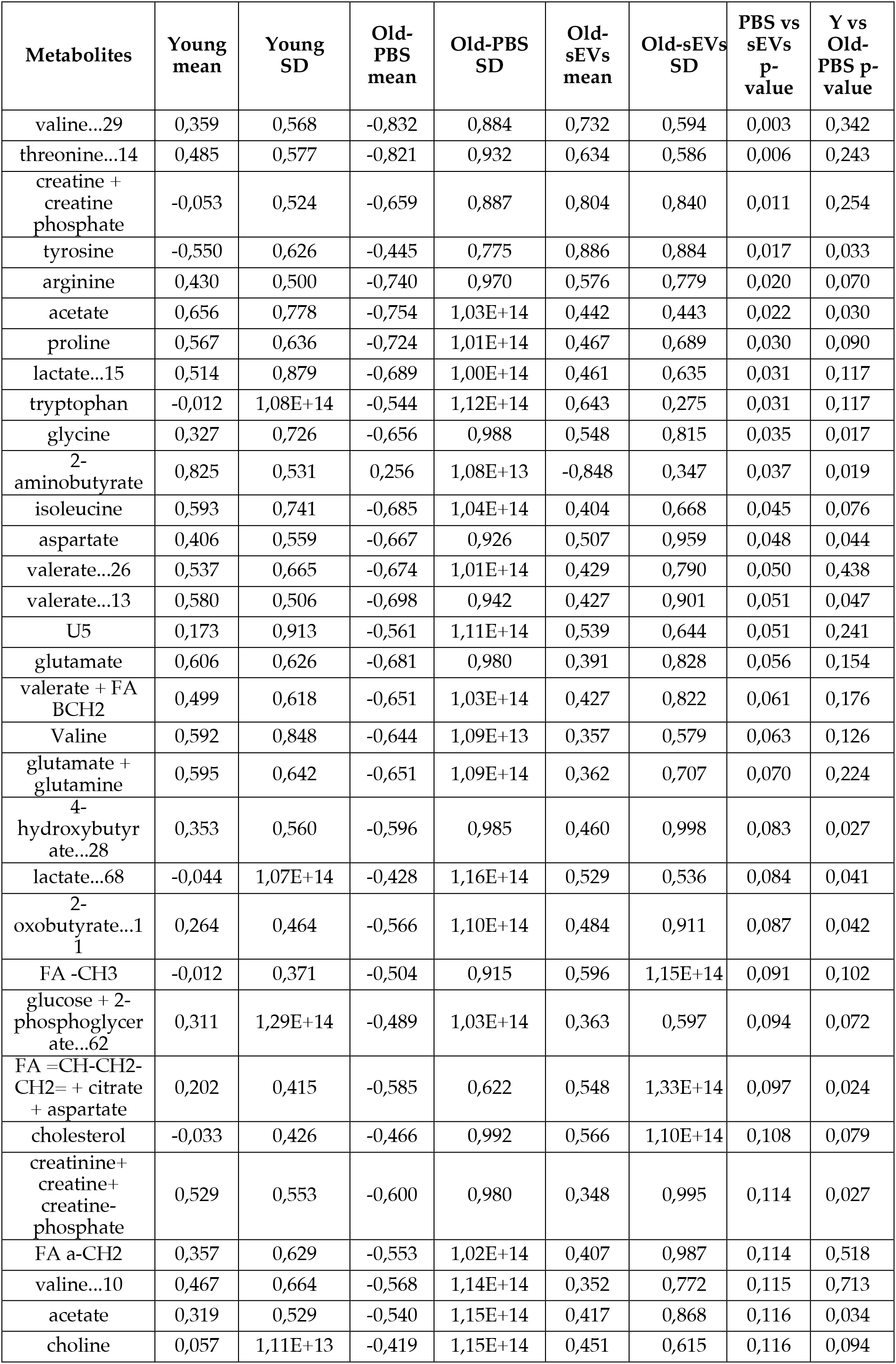
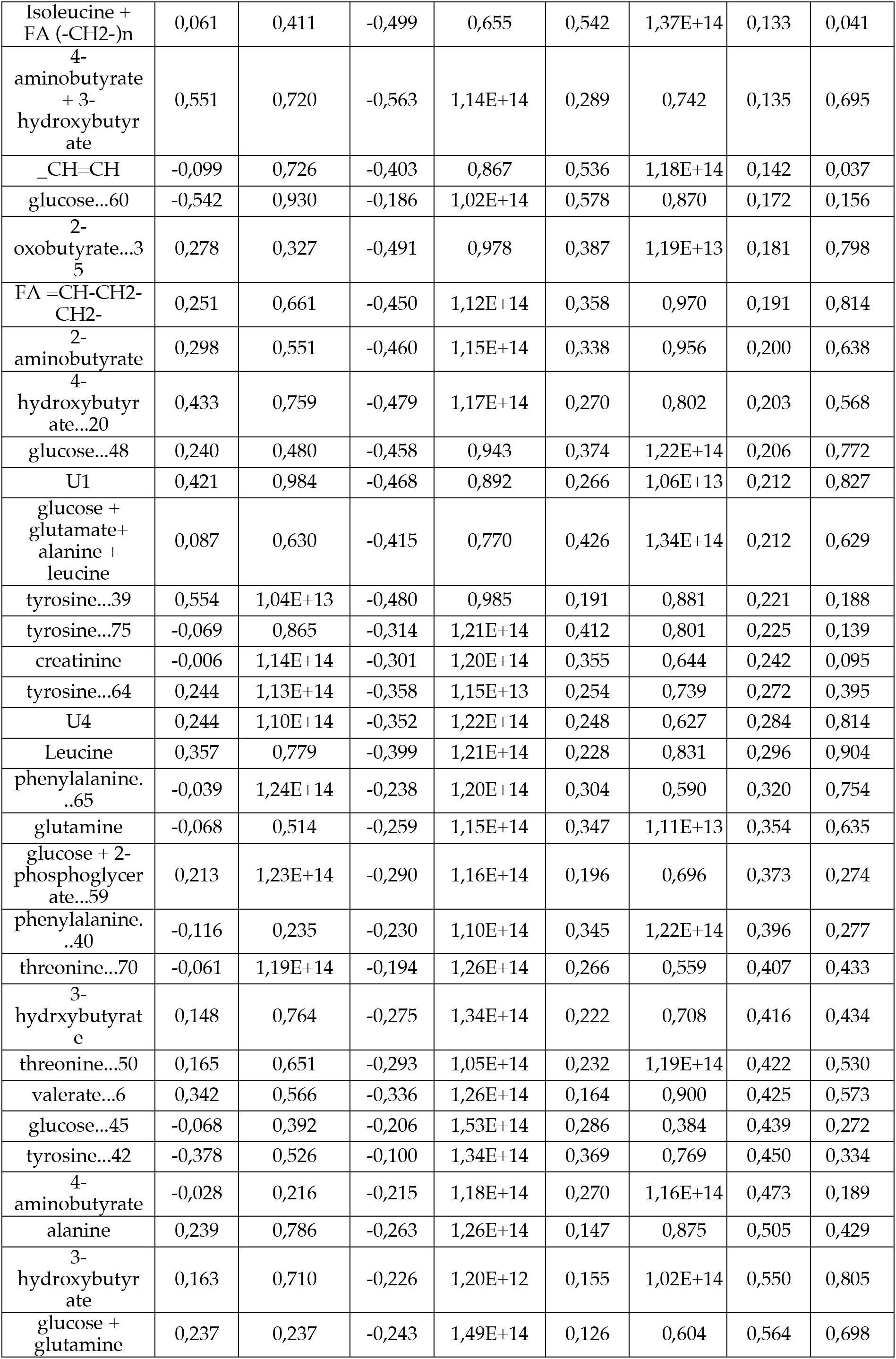
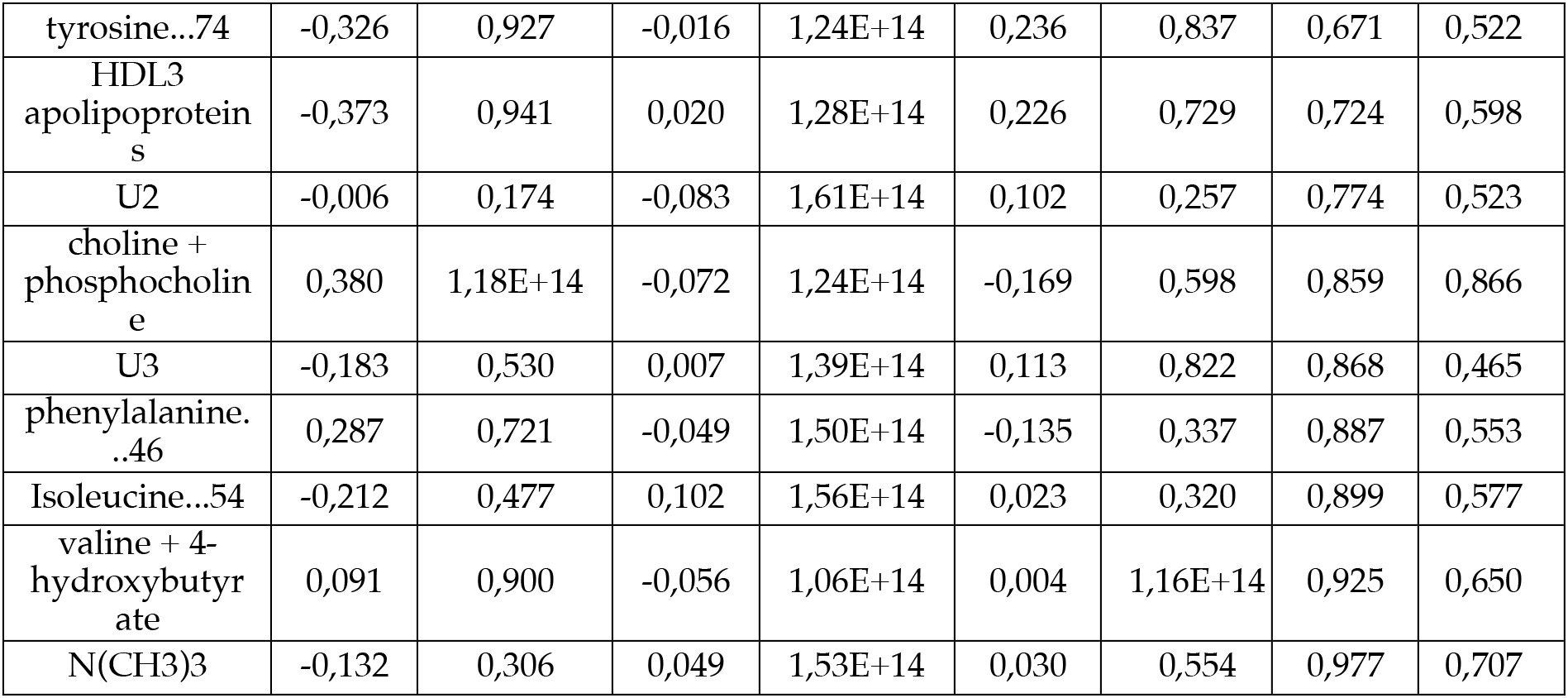
Data of the analysis of metabolites present in plasma comparing young mice, old mice treated with vehicle (PBS), and old mice treated with sEVs.

| Metabolites | Young mean | Young SD | Old-PBS mean | Old-PBS SD | Old-sEVs mean | Old-sEVs SD | PBS vs sEVs p-value | Y vs Old-PBS p-value |
| --- | --- | --- | --- | --- | --- | --- | --- | --- |
| valine...29 | 0,359 | 0,568 | -0,832 | 0,884 | 0,732 | 0,594 | 0,003 | 0,342 |
| threonine...14 | 0,485 | 0,577 | -0,821 | 0,932 | 0,634 | 0,586 | 0,006 | 0,243 |
| creatine + creatine phosphate | -0,053 | 0,524 | -0,659 | 0,887 | 0,804 | 0,840 | 0,011 | 0,254 |
| tyrosine | -0,550 | 0,626 | -0,445 | 0,775 | 0,886 | 0,884 | 0,017 | 0,033 |
| arginine | 0,430 | 0,500 | -0,740 | 0,970 | 0,576 | 0,779 | 0,020 | 0,070 |
| acetate | 0,656 | 0,778 | -0,754 | 1,03E+14 | 0,442 | 0,443 | 0,022 | 0,030 |
| proline | 0,567 | 0,636 | -0,724 | 1,01E+14 | 0,467 | 0,689 | 0,030 | 0,090 |
| lactate...15 | 0,514 | 0,879 | -0,689 | 1,00E+14 | 0,461 | 0,635 | 0,031 | 0,117 |
| tryptophan | -0,012 | 1,08E+14 | -0,544 | 1,12E+14 | 0,643 | 0,275 | 0,031 | 0,117 |
| glycine | 0,327 | 0,726 | -0,656 | 0,988 | 0,548 | 0,815 | 0,035 | 0,017 |
| 2-aminobutyrate | 0,825 | 0,531 | 0,256 | 1,08E+13 | -0,848 | 0,347 | 0,037 | 0,019 |
| isoleucine | 0,593 | 0,741 | -0,685 | 1,04E+14 | 0,404 | 0,668 | 0,045 | 0,076 |
| aspartate | 0,406 | 0,559 | -0,667 | 0,926 | 0,507 | 0,959 | 0,048 | 0,044 |
| valerate...26 | 0,537 | 0,665 | -0,674 | 1,01E+14 | 0,429 | 0,790 | 0,050 | 0,438 |
| valerate...13 | 0,580 | 0,506 | -0,698 | 0,942 | 0,427 | 0,901 | 0,051 | 0,047 |
| U5 | 0,173 | 0,913 | -0,561 | 1,11E+14 | 0,539 | 0,644 | 0,051 | 0,241 |
| glutamate | 0,606 | 0,626 | -0,681 | 0,980 | 0,391 | 0,828 | 0,056 | 0,154 |
| valerate + FA BCH2 | 0,499 | 0,618 | -0,651 | 1,03E+14 | 0,427 | 0,822 | 0,061 | 0,176 |
| Valine | 0,592 | 0,848 | -0,644 | 1,09E+13 | 0,357 | 0,579 | 0,063 | 0,126 |
| glutamate + glutamine | 0,595 | 0,642 | -0,651 | 1,09E+14 | 0,362 | 0,707 | 0,070 | 0,224 |
| 4-hydroxybutyrate...28 | 0,353 | 0,560 | -0,596 | 0,985 | 0,460 | 0,998 | 0,083 | 0,027 |
| lactate...68 | -0,044 | 1,07E+14 | -0,428 | 1,16E+14 | 0,529 | 0,536 | 0,084 | 0,041 |
| 2-oxobutyrate...11 | 0,264 | 0,464 | -0,566 | 1,10E+14 | 0,484 | 0,911 | 0,087 | 0,042 |
| FA -CH3 | -0,012 | 0,371 | -0,504 | 0,915 | 0,596 | 1,15E+14 | 0,091 | 0,102 |
| glucose + 2-phosphoglycerate...62 | 0,311 | 1,29E+14 | -0,489 | 1,03E+14 | 0,363 | 0,597 | 0,094 | 0,072 |
| FA =CH-CH2-CH2= + citrate + aspartate | 0,202 | 0,415 | -0,585 | 0,622 | 0,548 | 1,33E+14 | 0,097 | 0,024 |
| cholesterol | -0,033 | 0,426 | -0,466 | 0,992 | 0,566 | 1,10E+14 | 0,108 | 0,079 |
| creatinine+ creatine+ creatine-phosphate | 0,529 | 0,553 | -0,600 | 0,980 | 0,348 | 0,995 | 0,114 | 0,027 |
| FA a-CH2 | 0,357 | 0,629 | -0,553 | 1,02E+14 | 0,407 | 0,987 | 0,114 | 0,518 |
| valine...10 | 0,467 | 0,664 | -0,568 | 1,14E+14 | 0,352 | 0,772 | 0,115 | 0,713 |
| acetate | 0,319 | 0,529 | -0,540 | 1,15E+14 | 0,417 | 0,868 | 0,116 | 0,034 |
| choline | 0,057 | 1,11E+13 | -0,419 | 1,15E+14 | 0,451 | 0,615 | 0,116 | 0,094 |
| Isoleucine +<br>FA (-CH<sub>2</sub>)<sub>n</sub> | 0,061 | 0,411 | -0,499 | 0,655 | 0,542 | 1,37E+14 | 0,133 | 0,041 |
| 4-aminobutyrate<br>+ 3-hydroxybutyr<br>ate | 0,551 | 0,720 | -0,563 | 1,14E+14 | 0,289 | 0,742 | 0,135 | 0,695 |
| _CH=CH | -0,099 | 0,726 | -0,403 | 0,867 | 0,536 | 1,18E+14 | 0,142 | 0,037 |
| glucose...60 | -0,542 | 0,930 | -0,186 | 1,02E+14 | 0,578 | 0,870 | 0,172 | 0,156 |
| 2-oxobutyrate...3<br>5 | 0,278 | 0,327 | -0,491 | 0,978 | 0,387 | 1,19E+13 | 0,181 | 0,798 |
| FA =CH-CH<sub>2</sub>-<br>CH<sub>2</sub>- | 0,251 | 0,661 | -0,450 | 1,12E+14 | 0,358 | 0,970 | 0,191 | 0,814 |
| 2-aminobutyrate | 0,298 | 0,551 | -0,460 | 1,15E+14 | 0,338 | 0,956 | 0,200 | 0,638 |
| 4-hydroxybutyr<br>ate...20 | 0,433 | 0,759 | -0,479 | 1,17E+14 | 0,270 | 0,802 | 0,203 | 0,568 |
| glucose...48 | 0,240 | 0,480 | -0,458 | 0,943 | 0,374 | 1,22E+14 | 0,206 | 0,772 |
| U1 | 0,421 | 0,984 | -0,468 | 0,892 | 0,266 | 1,06E+13 | 0,212 | 0,827 |
| glucose +<br>glutamate+<br>alanine +<br>leucine | 0,087 | 0,630 | -0,415 | 0,770 | 0,426 | 1,34E+14 | 0,212 | 0,629 |
| tyrosine...39 | 0,554 | 1,04E+13 | -0,480 | 0,985 | 0,191 | 0,881 | 0,221 | 0,188 |
| tyrosine...75 | -0,069 | 0,865 | -0,314 | 1,21E+14 | 0,412 | 0,801 | 0,225 | 0,139 |
| creatinine | -0,006 | 1,14E+14 | -0,301 | 1,20E+14 | 0,355 | 0,644 | 0,242 | 0,095 |
| tyrosine...64 | 0,244 | 1,13E+14 | -0,358 | 1,15E+13 | 0,254 | 0,739 | 0,272 | 0,395 |
| U4 | 0,244 | 1,10E+14 | -0,352 | 1,22E+14 | 0,248 | 0,627 | 0,284 | 0,814 |
| Leucine | 0,357 | 0,779 | -0,399 | 1,21E+14 | 0,228 | 0,831 | 0,296 | 0,904 |
| phenylalanine.<br>..65 | -0,039 | 1,24E+14 | -0,238 | 1,20E+14 | 0,304 | 0,590 | 0,320 | 0,754 |
| glutamine | -0,068 | 0,514 | -0,259 | 1,15E+14 | 0,347 | 1,11E+13 | 0,354 | 0,635 |
| glucose + 2-phosphoglycer<br>ate...59 | 0,213 | 1,23E+14 | -0,290 | 1,16E+14 | 0,196 | 0,696 | 0,373 | 0,274 |
| phenylalanine.<br>..40 | -0,116 | 0,235 | -0,230 | 1,10E+14 | 0,345 | 1,22E+14 | 0,396 | 0,277 |
| threonine...70 | -0,061 | 1,19E+14 | -0,194 | 1,26E+14 | 0,266 | 0,559 | 0,407 | 0,433 |
| 3-hydrxybutyrat<br>e | 0,148 | 0,764 | -0,275 | 1,34E+14 | 0,222 | 0,708 | 0,416 | 0,434 |
| threonine...50 | 0,165 | 0,651 | -0,293 | 1,05E+14 | 0,232 | 1,19E+14 | 0,422 | 0,530 |
| valerate...6 | 0,342 | 0,566 | -0,336 | 1,26E+14 | 0,164 | 0,900 | 0,425 | 0,573 |
| glucose...45 | -0,068 | 0,392 | -0,206 | 1,53E+14 | 0,286 | 0,384 | 0,439 | 0,272 |
| tyrosine...42 | -0,378 | 0,526 | -0,100 | 1,34E+14 | 0,369 | 0,769 | 0,450 | 0,334 |
| 4-aminobutyrate | -0,028 | 0,216 | -0,215 | 1,18E+14 | 0,270 | 1,16E+14 | 0,473 | 0,189 |
| alanine | 0,239 | 0,786 | -0,263 | 1,26E+14 | 0,147 | 0,875 | 0,505 | 0,429 |
| 3-hydroxybutyr<br>ate | 0,163 | 0,710 | -0,226 | 1,20E+12 | 0,155 | 1,02E+14 | 0,550 | 0,805 |
| glucose +<br>glutamine | 0,237 | 0,237 | -0,243 | 1,49E+14 | 0,126 | 0,604 | 0,564 | 0,698 |
| tyrosine...74 | -0,326 | 0,927 | -0,016 | 1,24E+14 | 0,236 | 0,837 | 0,671 | 0,522 |
| HDL3<br>apolipoprotein<br>s | -0,373 | 0,941 | 0,020 | 1,28E+14 | 0,226 | 0,729 | 0,724 | 0,598 |
| U2 | -0,006 | 0,174 | -0,083 | 1,61E+14 | 0,102 | 0,257 | 0,774 | 0,523 |
| choline +<br>phosphocholin<br>e | 0,380 | 1,18E+14 | -0,072 | 1,24E+14 | -0,169 | 0,598 | 0,859 | 0,866 |
| U3 | -0,183 | 0,530 | 0,007 | 1,39E+14 | 0,113 | 0,822 | 0,868 | 0,465 |
| phenylalanine.<br>..46 | 0,287 | 0,721 | -0,049 | 1,50E+14 | -0,135 | 0,337 | 0,887 | 0,553 |
| Isoleucine...54 | -0,212 | 0,477 | 0,102 | 1,56E+14 | 0,023 | 0,320 | 0,899 | 0,577 |
| valine + 4-hydroxybutyr<br>ate | 0,091 | 0,900 | -0,056 | 1,06E+14 | 0,004 | 1,16E+14 | 0,925 | 0,650 |
| N(CH3)3 | -0,132 | 0,306 | 0,049 | 1,53E+14 | 0,030 | 0,554 | 0,977 | 0,707 |

**Supplementary Table 4:**
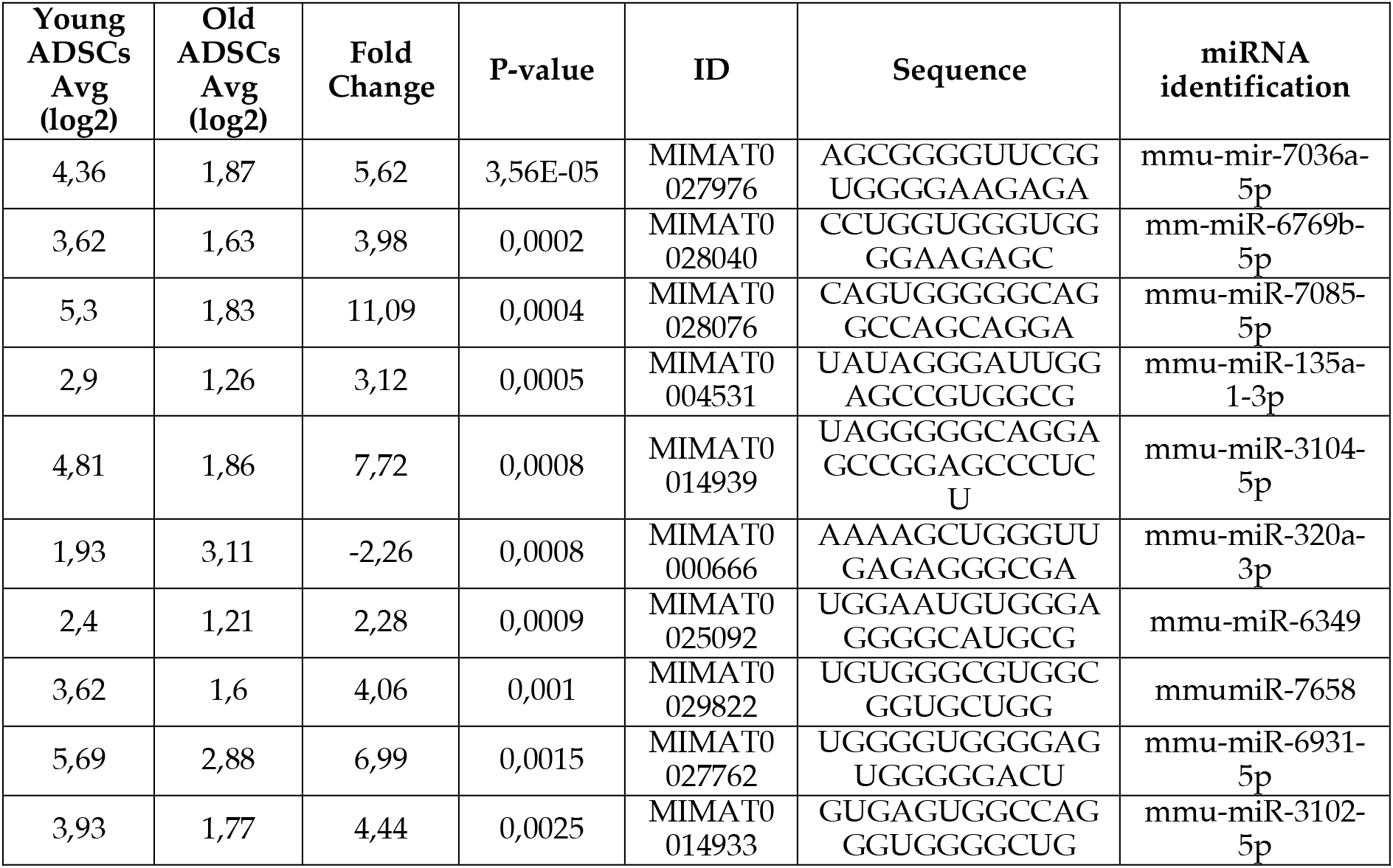
Data of the analysis of differential expressed miRNAs contained in sEVs from young ADSCs and old ADSCs.

**Supplementary Table 5:**
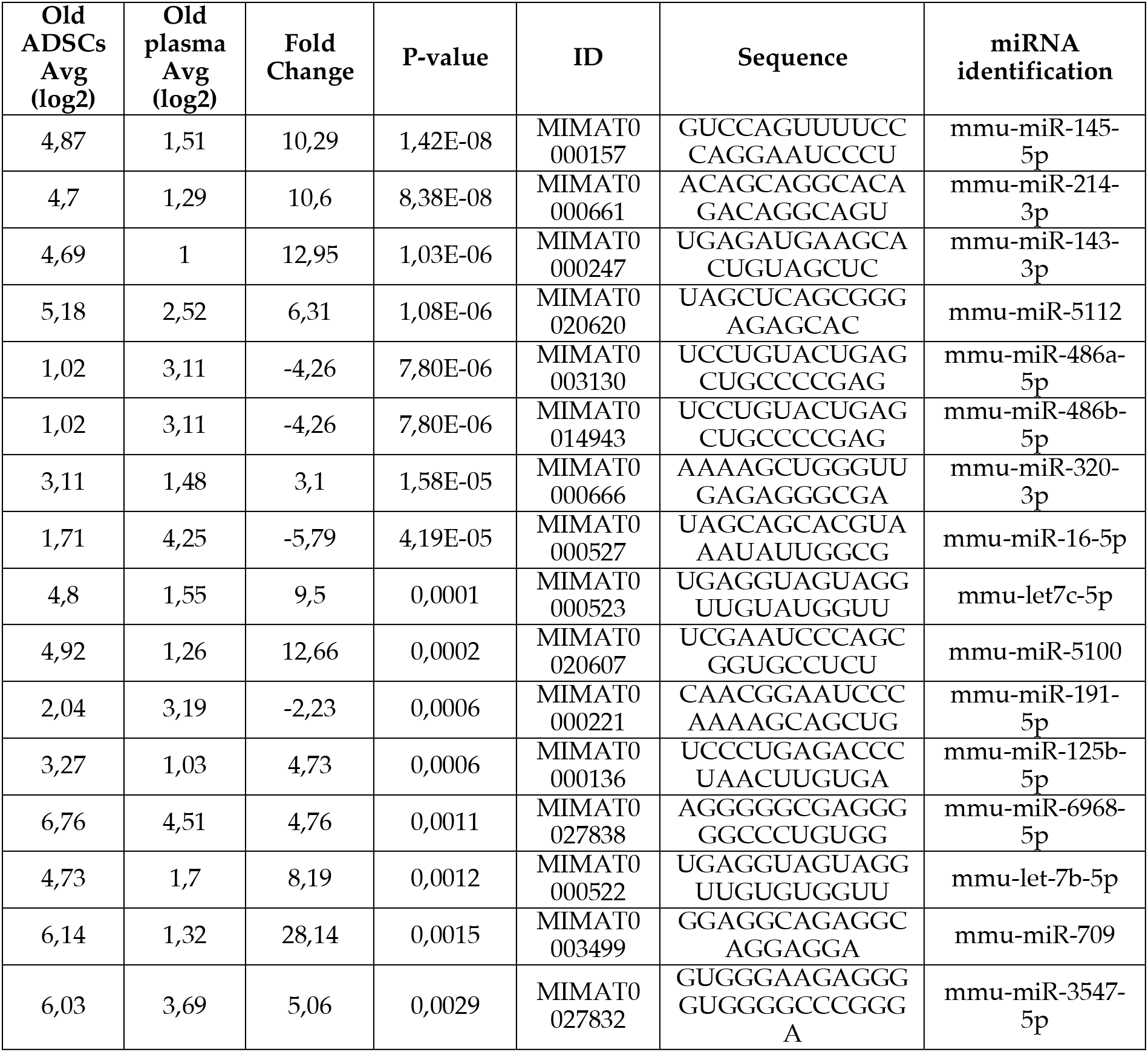
Data of the analysis of differential expressed miRNAs contained in sEVs from old ADSCs and plasma from aged mice

**Supplementary Table 6:**
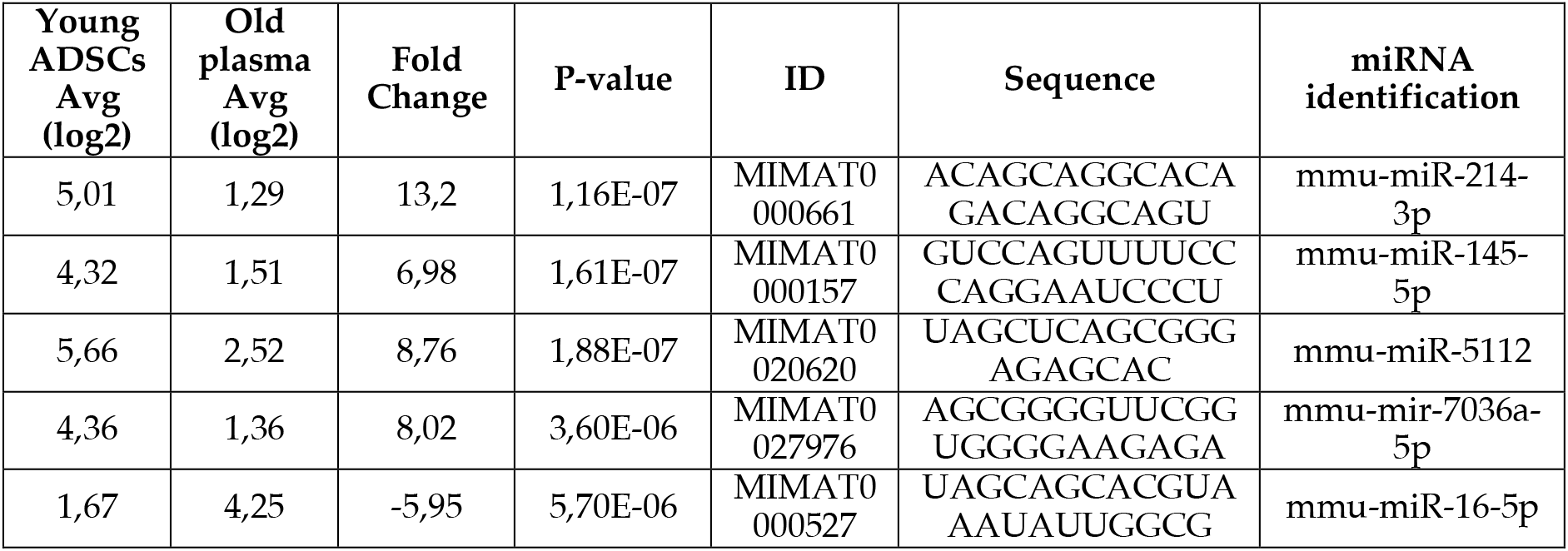
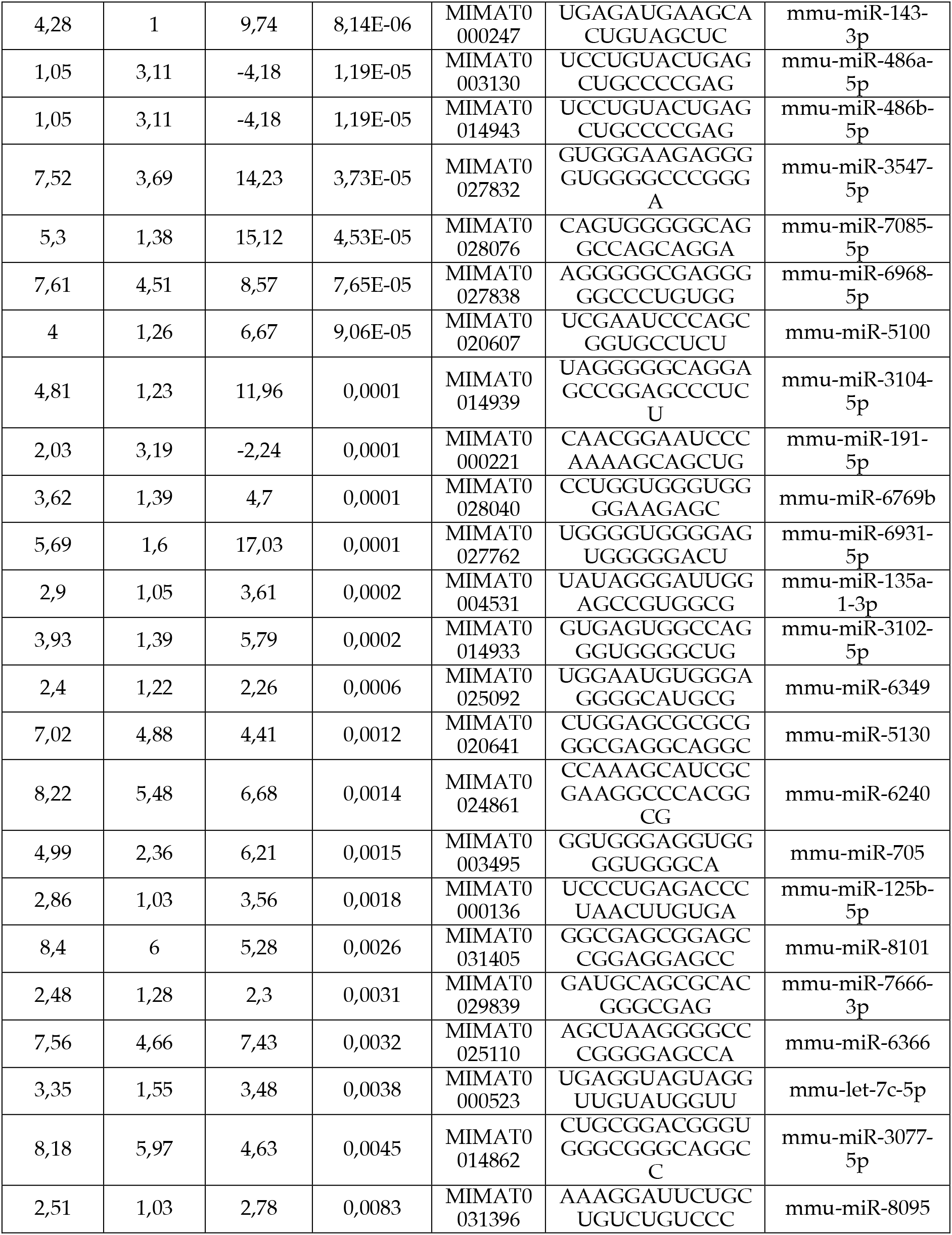
Data of the analysis of differential expressed miRNAs contained in sEVs from young ADSCs and plasma from aged mice.

